# Opposing actions of co-released GABA and neurotensin on the activity of preoptic neurons and on body temperature

**DOI:** 10.1101/2024.04.15.589556

**Authors:** Iustin V. Tabarean

## Abstract

Neurotensin (Nts) is a neuropeptide acting as a neuromodulator in the brain. Pharmacological studies have identified Nts as a potent hypothermic agent. The medial preoptic area, a region that plays an important role in the control of thermoregulation, contains a high density of neurotensinergic neurons and Nts receptors. The conditions in which neurotensinergic neurons play a role in thermoregulation are not known. In this study optogenetic stimulation of preoptic Nts neurons induced a small hyperthermia. *In vitro*, optogenetic stimulation of preoptic Nts neurons resulted in synaptic release of GABA and net inhibition of the preoptic pituitary adenylate cyclase-activating polypeptide (PACAP) neurons firing activity. GABA-A receptor antagonist or genetic deletion of VGAT in Nts neurons unmasked also an excitatory effect that was blocked by a Nts receptor 1 antagonist. Stimulation of preoptic Nts neurons lacking VGAT resulted in excitation of PACAP neurons and hypothermia. Mice lacking VGAT expression in Nts neurons presented changes in the fever response and in the responses to heat or cold exposure as well as an altered circadian rhythm of body temperature. Chemogenetic activation of all Nts neurons in the brain induced a 4-5 °C hypothermia, which could be blocked by Nts receptor antagonists in the preoptic area. Chemogenetic activation of preoptic neurotensinergic projections resulted in robust excitation of preoptic PACAP neurons. Taken together our data demonstrate that endogenously released Nts can induce potent hypothermia and that excitation of preoptic PACAP neurons is the cellular mechanism that triggers this response.

## Introduction

Homeotherms, including mammals, maintain core body temperature (CBT) within a narrow range, an essential requirement for survival. The preoptic area of the hypothalamus plays an important role in CBT regulation by integrating peripheral thermal information and by sending efferent signals that control thermal effector organs (1–3). Via projections to the dorsomedial hypothalamus and rostral raphe pallidus, preoptic neurons control thermogenesis as well as heat loss mechanisms (4). The preoptic area also orchestrates the fever response, an important mechanism for fighting infection or inflammatory disease (1, 5). Recent studies have identified populations of preoptic neurons that integrate peripheral thermal information and control the core body temperature (CBT) and play an important role in the fever response to endotoxin (6–9).

Neurotensin (Nts) is a 13 aminoacid peptide found in the central nervous system (CNS) as well as in the gastrointestinal tract (10). Nts-producing neurons and their projections are widely distributed in the brain, which may explain the wide variety of effects of this peptide (11). Pharmacological approaches have revealed a role of Nts in analgesia, arousal, blood pressure, feeding, reward, sleep, the stress response and thermoregulation (12–15). Recently, studies have started to unravel the role played by specific populations of neurotensinergic neurons in feeding (16, 17) and drinking behaviors (18) as well as in sleep (19, 20) and appetitive and reinforcing aspects of motivated behaviors (21).

When infused intracerebroventricularly or in the preoptic area Nts induces a 4-5 °C hypothermia (22–24). The effect was attributed to activation of heat loss mechanisms (25). The medial preoptic area (MPO) expresses high densities of both neurotensinergic neurons and neurotensin receptors (11, 26, 27). A previous study has identified a population of MPO Nts neurons that express VGAT and modulate sexual behavior via projections to the ventral tegmental area (21). The present study has investigated the cellular characteristics of MPO Nts neurons, their influence on the activity of nearby MPO neurons and on CBT as well as the effect of VGAT deletion on these actions.

## Results

### MPO^Nts;hChR2^ neurons are GABAergic and their optogenetic stimulation decreases the firing rate of MPO^PACAP^ neurons by increasing the frequency of IPSCs

Nts-cre mice received bilateral MPO injections of AAV5-EF1a-double floxed-hChR2(H134R)-EYFP-WPRE-HGHpA (see Methods) to express the excitatory opsin ChR2-EYFP in MPO Nts neurons. We refer to these mice as MPO^Nts;hChR2^. Patch-clamp recordings revealed that MPO^Nts;hChR2^ neurons were spontaneously active and the majority of them (16 out of 22 neurons studied) fired mostly doublets or triplets of action potentials (Fig 1A) while the rest fired single action potentials. The average firing rate at 36°C was 4.29±2.45 Hz (n=22). Intrinsic properties of PO/AH neurons were investigated using injections of square current pulses delivered in whole-cell configuration. The average capacitance of the neurons tested was 17.86±2.48 pF (n=22). All neurons tested displayed a low threshold spike (LTS) upon the end of a hyperpolarizing current injection (Fig 1B). Injection of positive currents generated bursts of doublets or triplets of action potentials (Fig 1B). To verify the presence of Nts transcripts in MPO^Nts;hChR2^ we have carried out scRT/PCR in these neurons. We could detect Nts transcripts in 7 out of 10 MPO^Nts;hChR2^ studied (Fig 1C). We have also tested the expression of the neuronal markers VGAT(Slc32a1) and Vglut2. VGAT was detected in 6 out of 7 Nts-expressing neurons (Fig 1C) while none of these cells expressed Vglut2 (Fig1 Suppl) suggesting that the majority of MPO^Nts;hChR2^ are GABAergic.

**Figure 1.**
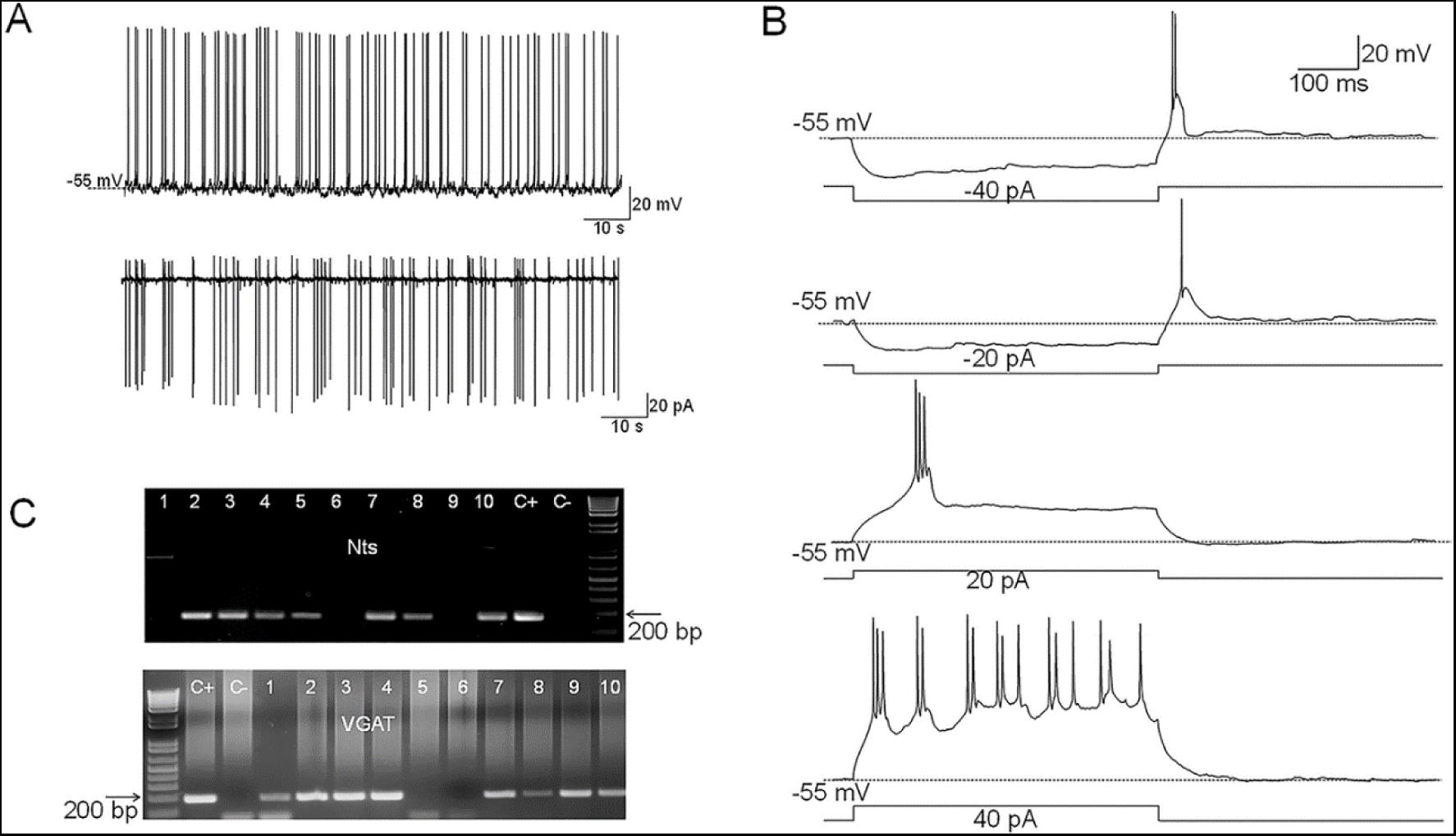
Electrophysiological characteristics of MPO^Nts;hChR2^ neurons. **A.** Representative example of spontaneous firing activity of MPO^Nts;hChR2^ neurons recorded in whole-cell (up) or cell-attached configuration (down). **B.** Membrane potential responses to hyperpolarizing current steps of -40 and -20 pA reveal the presence of a low threshold spike (LTS) upon depolarization to the resting membrane potential. Depolarizing current injections of 20 and 40 pA (right) elicit burst firing activity. The neuron fires 2-3 action potentials during each burst. **C.** Nts (up) and VGAT (down) transcripts are present in MPO^Nts;hChR2^ neurons. Representative results from 10 MPO^Nts;hChR2^ neurons. The expected sizes of the PCR product are 149 and 137 base pairs, respectively. Negative (−) control was amplified from a harvested cell without reverse-transcription, and positive control (+) was amplified using 1 ng of hypothalamic mRNA. Nts transcripts were detected in 7 out of 10 neurons while VGAT was detected in 8 neurons. 6 neurons expressed both transcripts.

Spot illumination of MPO^Nts;hChR2^ neurons instantly depolarized them and increased their firing rate in all neurons tested (not shown). Recordings from non-labeled MPO neurons revealed that spot illumination of nearby MPO^Nts;hChR2^ neurons resulted in a robust decrease in their firing rate (9 out of 30 neurons studied) or no effect in the others (21 out of 30 neurons studied). Fig 2A illustrates the decrease in firing rate of a MPO neuron (from 2.2 Hz to 0.9 Hz) induced by optogenetic stimulation of a nearby MPO^Nts;hChR2^ neuron. The effect was associated with a robust increase in the frequency of IPSPs from 0.5 Hz to 21.8 Hz (Fig 2A, inset). Overall, optogenetic stimulation of nearby MPO^Nts;hChR2^ neurons reduced the firing rate of the recorded MPO neurons from 5.10±1.32 Hz to 1.39±0.36 Hz (n=9). The control value for the firing rate was calculated as the average value during the 5 min period preceding the optogenetic stimulation. Upon ending the photostimulation a transient increase in firing rate was observed (Fig 2A). We therefore studied the effect of photostimulation during incubation with the GABA-A receptor antagonist gabazine (5 µM). In the presence of the antagonist the same neurons displayed the opposite effect, an increase in firing rate in response to photostimulation. Their firing rate was 7.89±2.61 Hz (n=9) in the presence of gabazine and increased to 9.57±2.83 Hz (n=9) during photostimulation (Fig 2A, B). Since we have previously found that exogenous Nts applied locally increased the firing activity of MPO neurons by activating NtsR1 (24) we have tested the effect of photostimulation in the presence of both gabazine (5 µM) and the NtsR1 antagonist SR48692 (100 nM). The antagonist blocked the effect of photostimulation indicating that the increase in firing rate depended on activation of NtsR1. MPO thermoregulatory neurons are a subgroup of the PACAP-expressing MPO population (7, 9). We have studied whether the MPO neurons excited by photostimulation (in the presence of gabazine) express PACAP transcripts. We have detected PACAP transcripts in 6 out of 8 neurons tested (Fig 2C).

**Figure 2.**
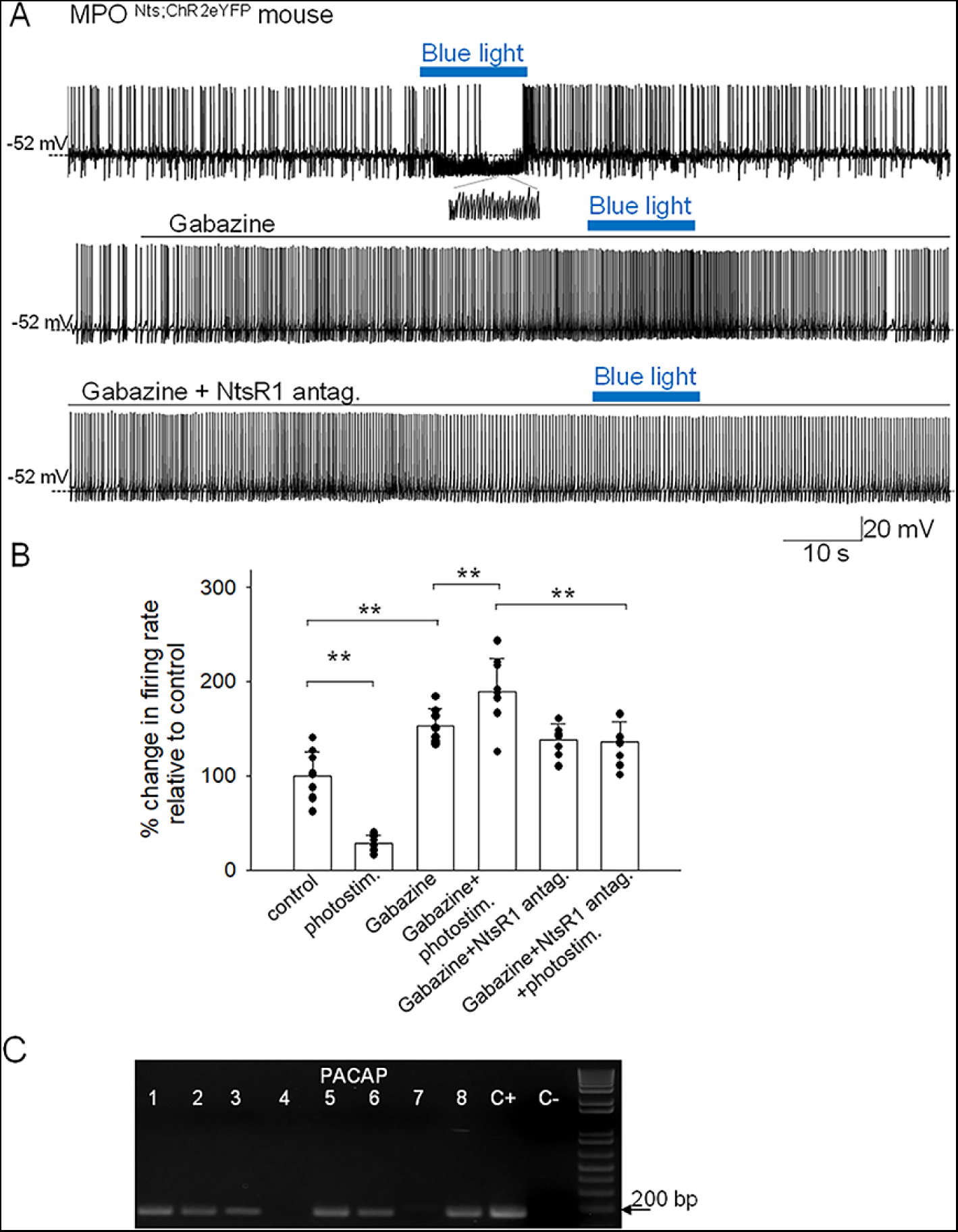
Effects of optogenetic stimulation of MPO^Nts;hChR2^ neurons on the firing activity of nearby MPO neurons. **A.** Optogenetic stimulation of a MPO^Nts;hChR2^ neuron decreases the spontaneous firing rate of a nearby MPO neuron (upper trace) from 2.2 Hz to 0.9 Hz and increases the frequency of IPSC from 0.5 Hz to 21.8 Hz (see expanded trace). Gabazine (5 µM) (middle trace) increased the spontaneous firing activity of the neuron and abolished the activation of IPSCs by optogenetic stimulation. In the presence of Gabazine optogenetic stimulation increased the firing activity of the neuron from 4.1 Hz to 9.6 Hz. This increase in firing activity was blocked by pe-incubation with the NtsR1 antagonist SR48692 (100 nM) (lower trace). **B.** Bar charts summarizing the effects of optogenetic stimulation of MPO^Nts;hChR2^ neurons on the spontaneous firing rates of nearby MPO neurons in control and in the presence of Gabazine (5 µM) and/or the NtsR1 antagonist SR48692 (100 nM). Bars represent means ± S.D. of the normalized firing rate relative to the control. The control value for the firing rate was calculated as the average value during the 5 min period preceding the optogenetic stimulation. Data pooled from n=9 neurons in each condition. There was a statistically significant difference between groups as determined by one-way ANOVA (F(5,40)= 77.71, p=1.08x10^-8^) followed by Tukey’s test between conditions; ** indicates statistical significance of P<0.01. The P-values of the Tukey’s statistical comparisons among groups are presented in Supplementary Table 2. **C.** PACAP transcripts are present in MPO neurons inhibited by optogenetic stimulation of nearby MPO^Nts;hChR2^ neurons. Representative results from 8 recorded MPO neurons. The expected size of the PCR product is 103 base pairs. Negative (−) control was amplified from a harvested cell without reverse-transcription, and positive control (+) was amplified using 1 ng of hypothalamic mRNA. PACAP transcripts were detected in 6 out of 8 neurons.

To further characterize the mechanisms involved in the modulation of the firing rates of MPO^PACAP^ neurons in response to optogenetic stimulation of MPO^Nts;hChR2^ we carried out additional experiments in voltage-clamp mode. In the presence of the glutamate receptor blockers CNQX (10 µM) and AP-5 (50 µM), as well as of the NtsR1antagonist SR48692 (100 nM), photostimulation MPO^Nts;hChR2^ neurons induced a robust increase in the IPSCs frequency (Fig 3A). These results indicate that GABA release was not dependent neither upon excitatory synaptic transmission nor upon activation of NtsR1, i.e. it is synaptically released by MPO^Nts;hChR2^ neurons. We have also found that brief photostimulation (10s or less) resulted in increased frequency of IPSCs only (Fig 3B, C) while longer photostimulation (30s or more) additionally activated an inward current (Fig 3B). The inward current was isolated in the presence of extracellular gabazine (5 µM) and averaged 9.26±2.39 pA (n=10). The inward current was abolished by bath application of NtsR1 antagonist SR48692 (100 nM) (Fig 3B, D). We have carried out a set of experiments in MPO^Nts;hChR2^ from female mice and recorded similar results. Optogenetic stimulation resulted in a robust increase in the frequency of IPSCs followed, with a delay of 20-40 s, by an inward current that averaged 11.3±3.4 pA (n=5) (Suppl Fig2). All the optogenetically-evoked responses were blocked by incubation with gabazine (5 µM) and SR48692 (100 nM) (Fig 3 Suppl).

**Figure 3.**
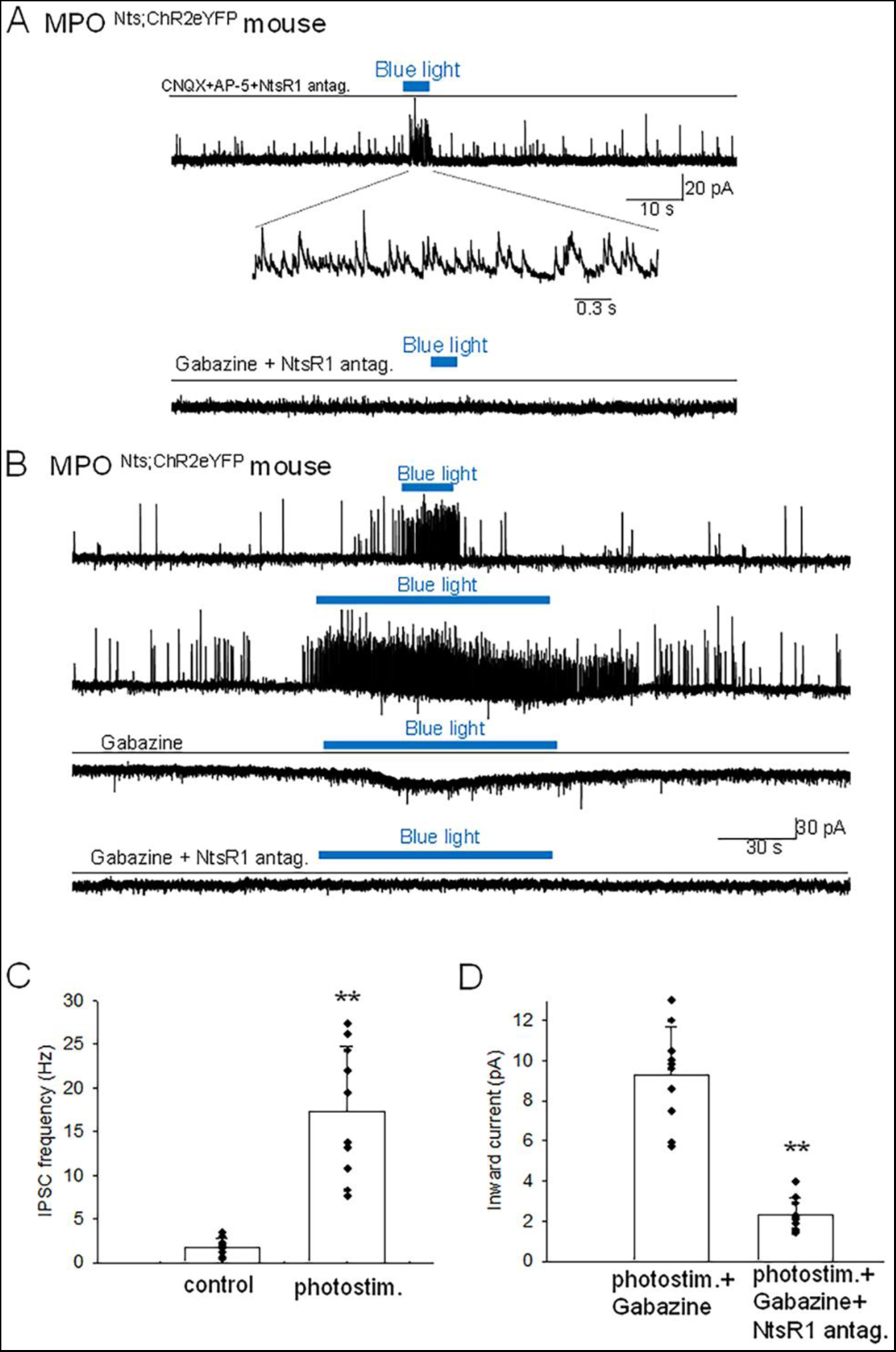
Optogenetic stimulation of MPO^Nts;hChR2^ neurons increases the frequency of IPSCs and activates an inward current in nearby MPO neurons. **A.** Optogenetic stimulation of a MPO^Nts;hChR2^ neuron activates IPSCs in a nearby MPO neuron. Recordings were performed in the presence of CNQX (20 µM), AP-5 (50 µM) and the NtsR1 antagonist SR48692 (100 nM). The sIPSCs were abolished by Gabazine (5 µM) (lower trace). The neuron was held at -50 mV. **B.** Optogenetic stimulation of the whole field of view containing several MPO^Nts;hChR2^ neurons for 20 s activates IPSCs in a nearby MPO neuron (upper trace). Longer optogenetic stimulation (80 s) of the same neurons activated both IPSCs and an inward current (middle traces). The inward current was abolished by the NtsR1 antagonist SR48692 (100 nM) (lower trace). The neuron was held at -50 mV. **C,D.** Bar charts summarizing the increase in the frequency of IPSCs (**C**) and the amplitude of the inward current (**D**) recorded in MPO neurons in response to optogenetic stimulation of several MPO^Nts;hChR2^ neurons. **C.** The IPSCs frequency increased from1.75±0.97 Hz to 17.8±7.47 Hz in response to photostimulation (one-way ANOVA (F(1,18)=42.5, p=4x10^-4^). The control value for the firing rate was calculated as the average value during the 5 min period preceding the optogenetic stimulation. **D**. The average inward current activated by optogenetic stimulation decreased from 9.26±2.39 pA to 2.31±0.83 pA in the presence of the NtsR1 antagonist SR48692 (100 nM) (one-way ANOVA (F(1,18)=75.35, p=7.5x10^-8^). Bars represent means ± S.D. Data pooled from n=10 neurons in each condition.

Finally, we have recorded from MPO^Nts;hChR2^ neurons and stimulated, using spot illumination, a different MPO^Nts;hChR2^ neuron to question the presence of reciprocal connections. None out of 8 MPO^Nts;hChR2^ neurons studied presented either IPSCs or an inward current evoked by optogenetic stimulation of other MPO^Nts;hChR2^ neurons.

Thermoregulatory preoptic neurons project to the dorsomedial hypothalamus, arcuate, paraventricular thalamus and to the rostral raphe pallidus to control thermoeffector mechanisms (8, 9, 28). We have examined the brains of MPO^Nts;eYFP^ mice in coronal slices to identify MPO projections to these sites, however no fluorescent signal was detected suggesting that MPO^Nts;eYFP^ modulate CBT by acting on MPO^PACAP^ neurons.

### Optogenetic stimulation of MPO^Nts;ChR2^ neurons *in vivo* induces hyperthermia. In contrast, optogenetic stimulation of MPO^Nts;VGAT-/-;ChR2^neurons induces hypothermia

Optogenetic activation of Nts neurons in the MPO using MPO^Nts;ChR2^ mice resulted in hyperthermia of up to 1.22±0.35 °C when compared with control MPO^Nts;eYFP^ mice (n=6 mice in each condition) (Fig 4A, B). To assess the role played by GABA release from MPO Nts neurons in the observed hyperthermia we activated Nts neurons in MPO^Nts;VGAT-/-;ChR2^ mice (see Methods). Surprisingly, a decrease in CBT of 1.44±0.29 °C was recorded relative to control MPO^Nts;VGAT-/-;eYFP^ mice (Fig 4C).

**Figure 4.**
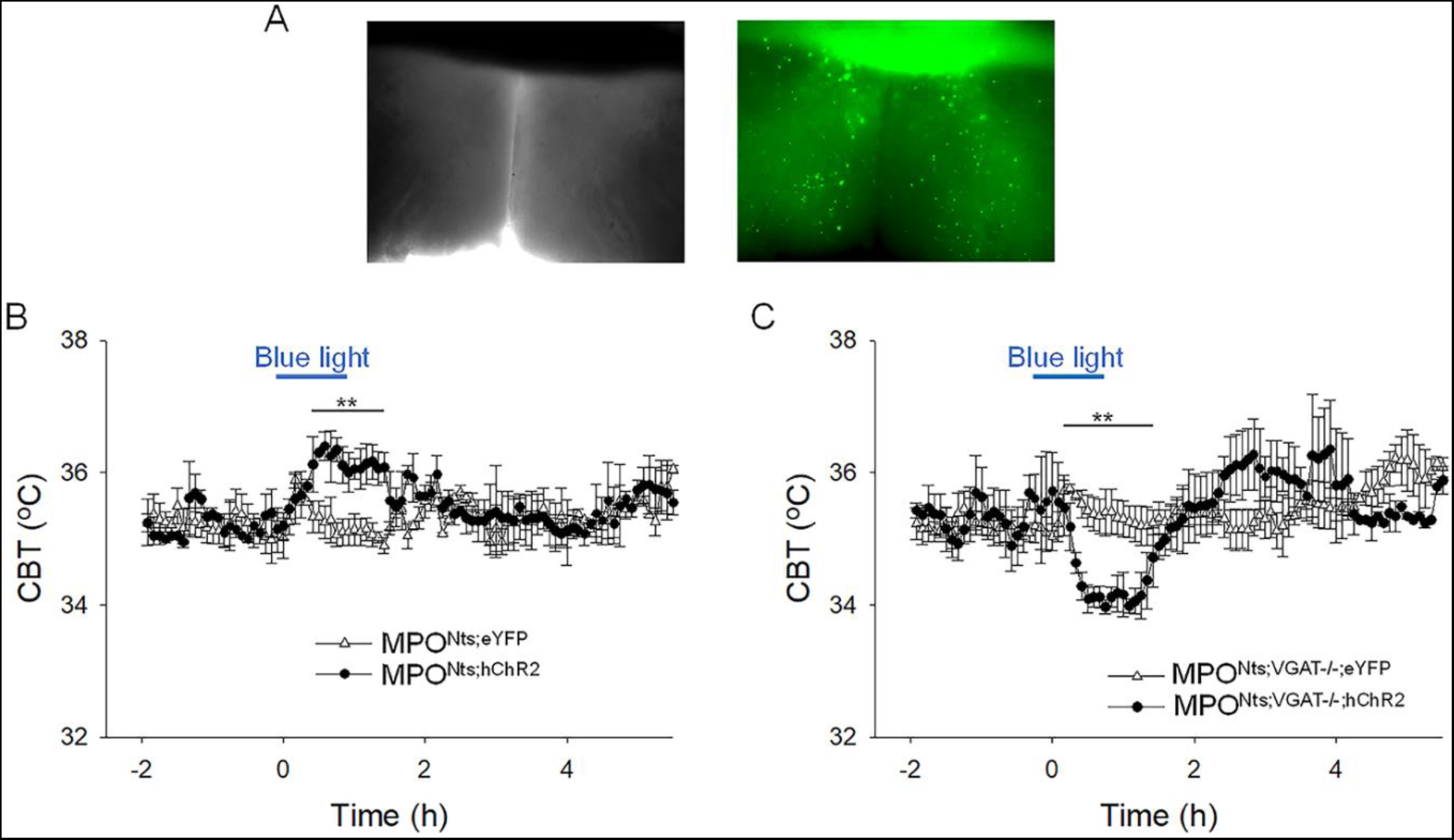
Optogenetic activation of MPO^Nts;hChR2^ neurons induces hyperthermia while optogenetic activation of MPO^Nts;VGAT-/-;hChR2^ neurons induces hypothermia. **A.** DIC (left) and fluorescence (right) images of an acute slice from MPO^Nts;hChR2^ mouse indicating hChR2-eYFP expression in the MPO. **B.** Optogenetic stimulation of MPO^Nts;hChR2^ neurons (●) *in vivo* for 1 hour (blue light) induced a hyperthermia of 1.22±0.35 °C relative to control (Δ). The response was statistically different to the response to photostimulation of control MPO^Nts;eYFP^ mice (Δ) (one-way repeated measures ANOVA, F(1,111)=20.9, p=1.2x10^-5^, followed by Man-Whitney U tests for each time point, ** P<0.01). **C.** Optogenetic stimulation of MPO^Nts;VGAT-/-;hChR2^ neurons (●) *in vivo* for 1 hour (blue light) induced a hypothermia of 1.44±0.29 °C relative to control (Δ). The response was statistically different to the response to photostimulation of control MPO^Nts;VGAT-/-;eYFP^ mice (Δ) (one-way repeated measures ANOVA, F(1,112)=8.27, p=4.8x10^-3^, followed by Man-Whitney U tests for each time point, ** P<0.01). **B,C.** The points represent averages±S.D. through the 7h recording period. Experiments were carried out in parallel in groups of 6 mice.

Similar results were obtained also in female MPO^Nts;ChR2^ mice and MPO^Nts;VGAT-/-;ChR2^ mice. Intra-MPO optogenetic stimulation resulted in a hyperthermia of 1.40±0.62 °C in MPO^Nts;ChR2^ females and in a hypothermia of 1.63±0.22 °C in MPO^Nts;VGAT-/-;ChR2^ females (Fig4 Suppl).

### Optogenetic stimulation of MPO^Nts; VGAT-/-;ChR2^ neurons increases the firing rate of MPO^PACAP^ neurons by activating an inward current

The basal electrophysiological properties of MPO^Nts; VGAT-/-;ChR2^ neurons were similar with those of MPO^Nts;ChR2^ neurons. The neurons were spontaneously active, with an average firing rate at 36 °C of 4.79±2.45 Hz (n=26). Recordings from non-labeled MPO neurons revealed that spot illumination of nearby MPO^Nts; VGAT-/-;ChR2^ neurons increased their firing rate (12 out of 26 neurons studied) or had no effect in the others (14 out of 26 neurons). Fig 5A illustrates the increase in firing rate of a MPO neuron (from 1.6 Hz to 3.9 Hz) induced by optogenetic stimulation of a nearby MPO^Nts;ChR2^ neuron, an effect which was associated with an apparent depolarization. Overall, optogenetic stimulation of nearby MPO^Nts;VGAT-/-;ChR2^ neurons increased the firing rate of the recorded MPO neurons from 5.68±1.66 Hz to 12.20±3.65 Hz (n=12) (Fig 5B). This effect was blocked by preincubation with the NtsR1 antagonist SR48692 (100 nM) (Fig 5A,B). In voltage-clamp experiments photostimulation activated an inward current which was abolished by bath application of NtsR1 antagonist SR48692 (100 nM) (Fig 5C, D). Using s.c. RT/PCR assays we have detected PACAP transcripts in 8 out of 11 neurons excited by photostimulation tested.

**Figure 5.**
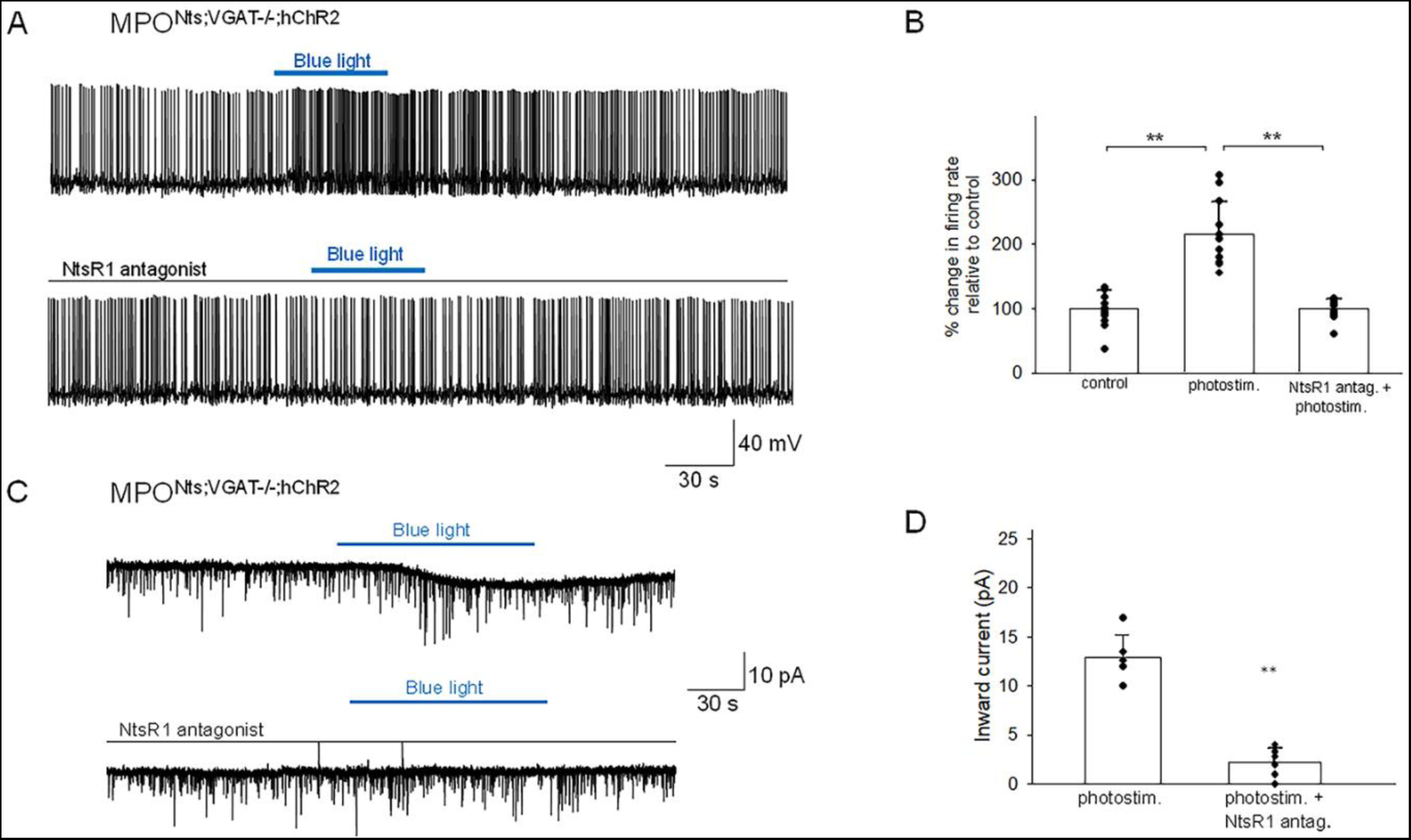
Optogenetic stimulation of MPO^Nts-VGAT-/-;hChR2^ neurons increases the firing activity of nearby MPO neurons. **A.** Optogenetic stimulation of MPO^Nts;VGAT-/-;hChR2^ neurons increases the spontaneous firing rate of a nearby MPO neuron (upper trace) from 1.6 Hz to 3.9 Hz. The photostimulation-induced increase in firing activity was blocked by pe-incubation with the NtsR1 antagonist SR48692 (100 nM) (lower trace). **B.** Bar charts summarizing the effects of optogenetic stimulation of MPO^Nts;VGAT-/-;hChR2^ neurons on the spontaneous firing rates of nearby MPO neurons in control and in the presence of the NtsR1 antagonist SR48692 (100 nM). Bars represent means ± S.D. of the normalized firing rate relative to the control. Data pooled from n=12 neurons in each condition. There was a statistically significant difference between groups as determined by one-way ANOVA (F(2,22)= 42.45, p=2.80x10^-8^) followed by Tukey’s test between conditions; ** indicates statistical significance of P<0.01, * indicates P<0.05. The P-values of the Tukey’s statistical comparisons among groups are presented in Supplementary Table 3. **C.** Optogenetic stimulation of the whole field of view containing several MPO^Nts;hChR2^ neurons for 20 s activates IPSCs in a nearby MPO neuron (upper trace). Longer optogenetic stimulation (80 s) of the same neurons activated both IPSCs and an inward current (middle traces). The inward current was abolished by the NTSR1 antagonist SR48692 (100 nM) (lower trace). The neuron was held at -50 mV. **D.** Bar chart summarizing the amplitude of the inward current recorded in MPO neurons in response to optogenetic stimulation in control and during incubation with the NtsR1 antagonist SR48692 (100 nM). The average inward current activated by optogenetic stimulation decreased from 12.86±2.33 pA to 2.18±1.49 pA in the presence of MPO^Nts;VGAT-/-;hChR2^ neurons the NtsR1 antagonist SR48692 (100 nM) (one-way ANOVA (F(1,22)=193.73, P= 2.49x10^-8^). Bars represent means ± S.D. Data pooled from n=12 neurons.

We have studied the presence of VGAT and Nts transcripts in MPO slices from Nts^VGAT-/-^ and in control Nts^VGAT+/+^ mice (Fig 5 Suppl). Nts and VGAT transcripts overlapped in control tissue while there was no colocalization in tissue from Nts^VGAT-/-^ mice (Fig 5 Suppl).

### Characterization of the fever response, the heat exposure and the cold exposure responses as well as of the circadian CBT profile in Nts^VGAT-/-^mice

Since MPO^Nts; VGAT-/-;ChR2^ mice displayed drastically different CBT responses to optogenetic stimulation relative to those of MPO^Nts;ChR2^ mice, suggesting that GABA release by Nts neurons played an important role in thermoregulation, we decided to further characterize thermoregulatory responses in the Nts^VGAT-/-^mice.

The LPS-induced fever was significantly different in Nts^VGAT-/-^ mice when compared with that of wild-type littermates Nts^VGAT+/+^ (Fig 6A). During the intermediate and late phases of fever the CBT was significantly lower in Nts^VGAT-/-^mice (Fig 6A).

**Figure 6.**
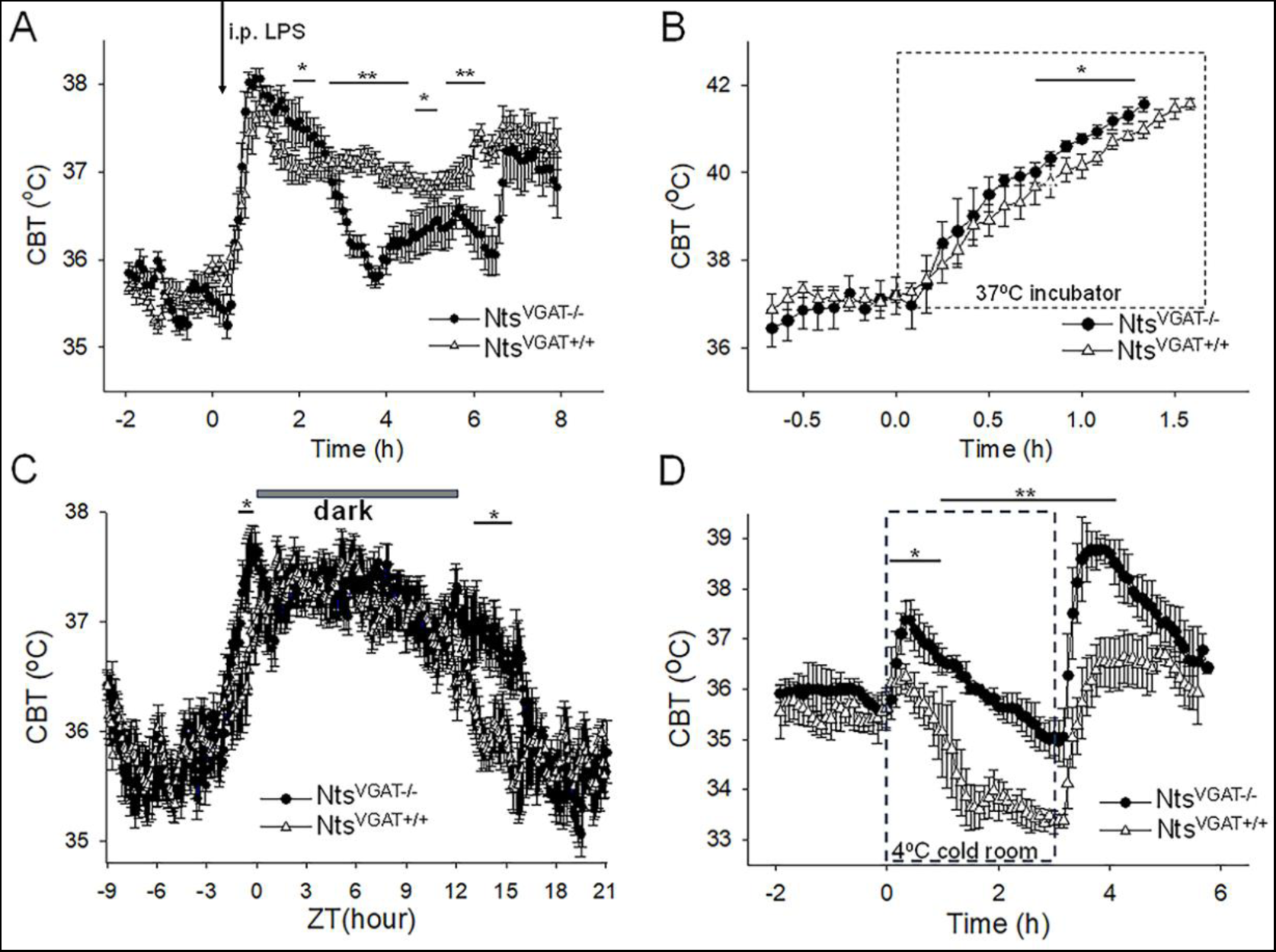
Altered thermoregulatory profile of Nts^VGAT-/-^ mice. **A.** CBT responses to i.p. injection (arrow) of LPS (0.03 mg/kg) in Nts^VGAT+/+^ mice (Δ, control) and Nts^VGAT-/-^ mice (●). LPS induced fever responses with differential profiles (one-way repeated measures ANOVA, F(1,94)=9.31, P=2.948x10^-3^, followed by Man-Whitney U tests for each time point, ** P<0.01, * P<0.05). **B.** CBT responses during a heat test in an incubator at 37 °C. The CBT increased faster in Nts^VGAT-/-^ mice (●) than in Nts^VGAT+/+^ mice (Δ, control) (repeated measures ANOVA, F(1,16)= 33.49, p= 2.78x10^-5^, followed by Man-Whitney U tests for each time point, * P<0.05). **C.** Circadian CBT profiles in Nts^VGAT-/-^ mice (●) than in Nts^VGAT+/+^ mice (Δ, control). Nts^VGAT-/-^ mice display a longer active phase relative to controls (repeated measures ANOVA, F(1,359)= 173.35, P=1.10x10^-9^, followed by Man-Whitney U tests for each time point, * P<0.05). Data for each mouse represents the average of 10 different 24 hour periods. **D.** CBT responses during a cold test in an incubator at 4°C. Nts^VGAT-/-^ mice (●), in contrast with Nts^VGAT+/+^ mice (Δ, control), displayed a significant hyperthermia at the beginning as well as following the end of the cold exposure (repeated measures ANOVA, F(1,92)=220.89, p= 4.3x10^-14^, followed by Man-Whitney U tests for each time point, * P<0.05, ** P<0.01). **A-D.** The points represent averages±S.D. (n= 6 male mice).

During heat exposure the CBT increased at a faster rate in Nts^VGAT-/-^ mice than in Nts^VGAT+/+^ mice (Fig 6B). The Nts^VGAT-/-^ mice and Nts^VGAT+/+^ mice reached 41.5 °C within 1.33±0.04 h and 1.53±0.04 h, respectively (Kruskal-Wallis, P=2.14x10^-4^, n=9 mice in each group). The Nts^VGAT-/-^ mice displayed significantly higher CBT (by ∼0.5 °C) after 45 min of heat exposure (Fig 6B).

The circadian CBT profile of Nts^VGAT-/-^ mice was also altered relative to that of Nts^VGAT+/+^ mice: the active phase started ∼1h earlier and ended ∼3h later (Fig 6C).

During a 3h exposure in a cold room at 4 °C the two mouse lines displayed strikingly different CBT responses (Fig 6D). The control mice displayed a gradual decrease in CBT which stabilized at 33-34 °C by third hour of the cold exposure. In contrast, Nts^VGAT-/-^ mice displayed a hyperthermia of 37.4 °C at the beginning of the experiment followed by a slow decline that reached ∼35 °C at the end of the three hours. Upon return to vivarium the CBT of controls returned to 37 °C while the Nts^VGAT-/-^mice displayed a transient hyperthermia to 39 °C.

### Activation of central Nts^hM3D(Gq)^ neurons induces potent hypothermia

Previous studies have reported the intracerebroventricular or intra MPO infusions of exogenous Nts induce potent hypothermia (23, 24). Since our results indicated that activation of MPO Nts neuron does not mimic this effect we hypothesized that Nts release by Nts neurons in other brain regions may be required to induce hypothermia. In order to activate all central Nts neurons we have generated Nts^hM3D(Gq)^ mice (see Methods) and employed chemogenetic stimulation. Activation of central Nts^hM3D(Gq)^ neurons induced a potent hypothermia of 4.8±0.6 °C relative to Nts-cre mice (control) (Fig 7A), value comparable with that obtained with MPO infusions of exogenous Nts (24). To test the hypothesis that the main locus of action is the MPO and to determine the possible role of NtsR1 and NtsR2 in this effect we have performed the chemogenetic activation of Nts^hM3D(Gq)^ in the presence of NtsR1 and/or NtsR2 antagonists in the MPO (Fig7B). The antagonists (or aCSF as control) were infused via a bilateral guide cannula 1.5 hours prior to CNO injection (i.p. 1 mg/kg). The NtsR1 antagonist SR48692 (300 nM, 100nl, blue trace) decreased the chemogenetically induced hypothermia by 41 % relative to control (Fig 7B). Higher concentrations of SR48692 (600 nM and 1.5 μM) resulted in similar reductions in the hypothermic response of 43 and 31%, respectively (n=6 mice in each condition). Following intra-MPO injections of NTSR2 antagonist NTRC 824 (200 nM, 100nl, black trace) the CNO-induced hypothermia was decreased by 32% relative to control (Fig 7B). When the two doses of antagonists were infused together the chemogenetically-induced hypothermia decreased by 76% (Fig 7B, red trace). These results indicate that the MPO accounts for most of the hypothermia induced by Nts^hM3D(Gq)^ neurons and that both receptor subtypes play an important role.

**Figure 7.**
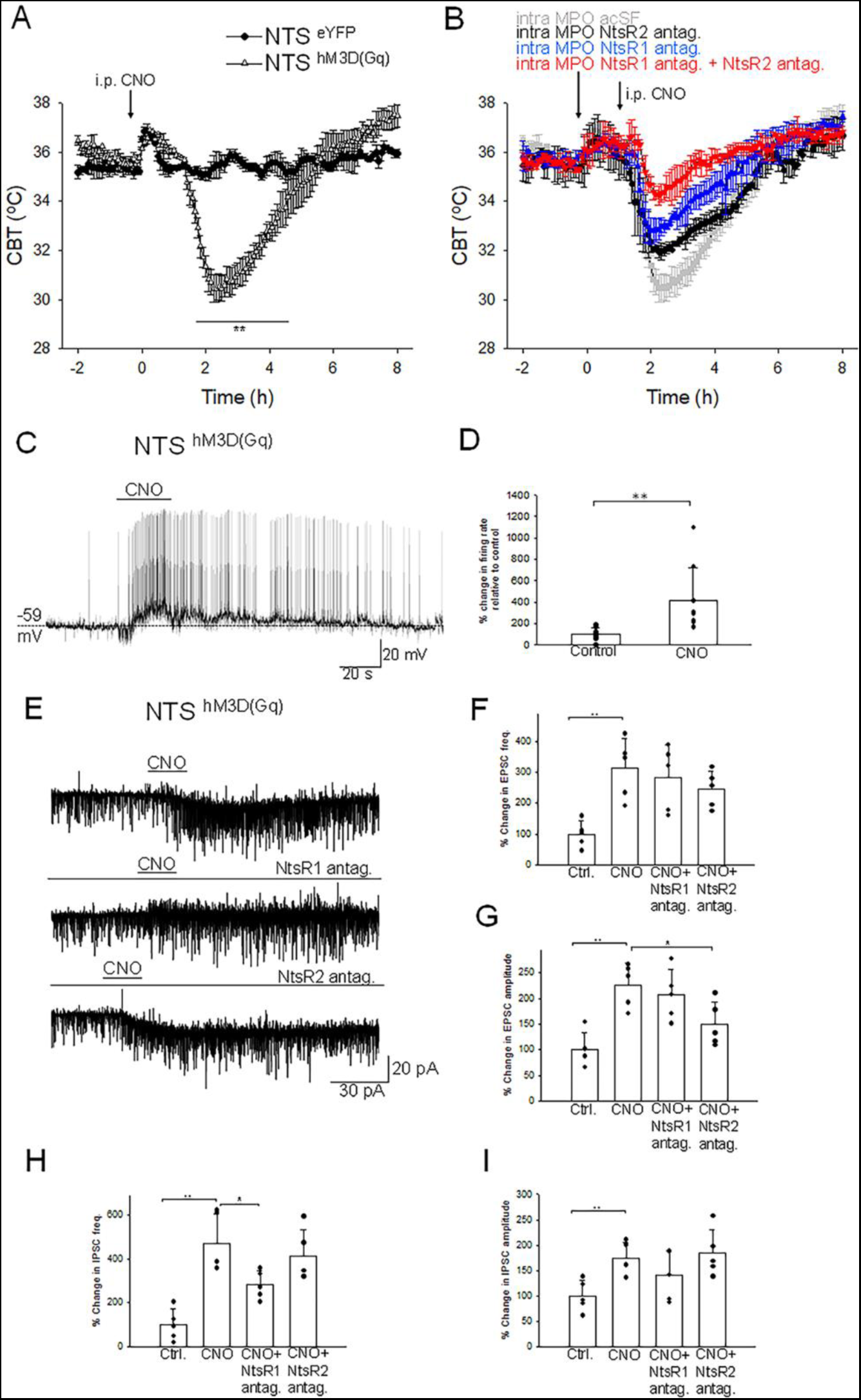
Chemogenetic activation of neurotensinergic neurons and projections induces hypothermia and potently excites MPO neurons. **A.** I.p. injection (arrow) of CNO (20 mM, 3 µl) in Nts^hM3D(Gq)^ mice (Δ) and in Nts-cre mice (control, ●). CNO induced a hypothermia of 4.8±0.6 °C (repeated measures ANOVA, F(1,242)=21.72, p=2.8x10^-6^, followed by followed by Man-Whitney U tests for each time point, ** P<0.01). **B.** Role of NtsR1 and NtsR2 expressed in the MPO in the CNO-induced activation of Nts^hM3D(Gq)^ neurons. Nts^hM3D(Gq)^ mice received a bilateral infusion of aCSF (gray), NtsR1 antagonist SR48692 (300 nM, 100nl, blue), NtsR2 antagonist NTRC 824 (200 nM, 100nl, black) and NtsR1 antagonist (300 nM, 100nl) + NtsR2 antagonist (200 nM, 100nl) (red) 1.5h prior to an i.p. injection of CNO (20 mM, 3 µl). The antagonists significantly reduced the hypothermia (repeated measures ANOVA, F(3,360)=71.33, p=8.82x10^-16^). **A,B.** The points represent averages±S.D. Experiments were carried out in parallel in groups of 6 **C.** Chemogenetic activation of neurotensinergic neurons and neurotensinergic projections in the MPO by bath application of CNO (3 µM) in slices from Nts^hM3D(Gq)^ mice depolarizes and increases the firing rate of a MPO neuron. The firing rate increased from 0.15 Hz to 1.65 Hz. **D.** Bar chart summarizing the effect of chemogenetic activation of neurotensinergic neurons and projections in the MPO on the firing rates of MPO neurons. There was a statistically significant difference between groups as determined by one-way ANOVA (F(1,16)=8.99, P=8.52x10^-3^; ** indicates statistical significance of P<0.01). Bars represent means ± S.D. of the normalized firing rate relative to the control. Data pooled from n=10 neurons. **E.** Chemogenetic activation of neurotensinergic neurons and neurotensinergic projections in the MPO by bath application of CNO (3 µM) in slices from Nts^hM3D(Gq)^ mice increases the amplitudes and frequencies of both IPSCs and EPSCs and activates an inward current (upper trace). The NtsR1 antagonist SR48692 (100 nM) abolished the inward current activated by CNO (middle trace). The NtsR2 antagonist NTRC 824 (100 nM) did not change the inward current activated by CNO but significantly decreased the amplitude of sEPSCs (lower trace). The neuron was held at -50 mV. **F,G,H,I.** Bar charts summarizing the increase in sEPSCs frequency (**F**) and amplitude (**G**) and IPSCs frequency (**H**) and amplitude (**I**). Bars represent means ± S.D. of the normalized frequency relative to the control. Data pooled from n=6 neurons. The changes were statistically significant for sEPSCs frequency (one-way ANOVA (F(3,19)=6.94, P=3.33x10^-3^) and amplitude (one-way ANOVA (F(3,19)=9.20, P=9.03x10^-4^) as well as for sIPSCs frequency (one-way ANOVA (F(3,19)=12.61, P=1.71x10^-4^) and amplitude (one-way ANOVA (F(3,19)=4.53, P=1.76x10^-3^). The P values for the inter-group comparisons are listed in Supplementary Tables 4-7.

### Chemogenetic activation of NTS^hM3D(Gq)^ neurons and projections in the MPO results in potent excitation of MPO^PACAP^

In order to understand the mechanisms by which chemogenetic activation of NTS^hM3D(Gq)^ neurons induced hypothermia, we have studied the effect of CNO on the activity of MPO non-labeled neurons in acute slices from NTS^hM3D(Gq)^ mice. Bath application of CNO excited 9 out 20 neurons studied and had no effect in the others. Fig 7C illustrates such a response in a non-labeled MPO neuron. Interestingly prior to depolarization an increase in the frequency of sIPSPs (arrow) was recorded. The average firing rate increased from 4.15±2.95 Hz to 10.63±5.83 Hz (n=9, ANOVA, F(1/16)=8.99 P=8.52x10^-3^)) in response to CNO (3 µM) (Fig 7D). S.c. RT/PCR assays identified PACAP transcripts in 7 of the 9 neurons excited by CNO tested. In voltage-clamp experiments bath application of CNO (3 µM) activated an inward current and potently increased the frequencies and amplitudes of both sEPSCs and sIPSCs (Fig7 E). The frequencies of sEPSCs and sIPSCs increased from 2.76±1.16 Hz to 9.14±5.36 Hz and 1.98±1.43 Hz to 8.84±6.38 Hz, respectively (n=6)(Fig 7F,H). The amplitudes of sIPSCs and sEPSCs increased from 13.61±4.51 pA to 29.60±5.32 pA and 13.02±3.99 pA to 22.12±5.53 pA, respectively (n=6)(Fig 7G,I). The NtsR1 antagonist SR48692 (100 nM) abolished the inward current activated by CNO (Fig 7E, middle trace) and significantly reduced the effect on the frequency of sIPSCs (Fig 7H). The NtsR2 antagonist NTRC 824 (100 nM) did not change the inward current activated by CNO but significantly decreased the amplitude of sEPSCs (Fig 7G).

## Discussion

In this study we have characterized a novel population of MPO neurons that express Nts (MPO^Nts^). These neurons are GABAergic as indicated by detection of VGAT transcripts as well as by the activation of gabazine-sensitive IPSCs in postsynaptic neurons in response to spot illumination of MPO^Nts;hChR2^ neurons. Interestingly, this population had a tendency to burst in response to depolarization, a characteristic of neurosecretory neurons (29, 30). In response to optogenetic stimulation MPO^Nts;hChR2^ neurons released both GABA and Nts as indicated by pharmacological experiments. Postsynaptically, increased frequency of IPSCs was detected instantaneously upon photostimulation while an inward current was activated after 20-40 seconds, indicating a differential timecourse of action. In view of its slower timecourse of action, Nts may act to limit and/or shorten the inhibitory effect of GABA on the activity of the postsynaptic neurons following increased burst of activity of MPO^Nts^ neurons. Nevertheless, the net effect during photostimulation was a decrease in the firing rate of the postsynaptic neuron, an increase in firing rate being unmasked only upon ending photostimulation or in the presence of gabazine. Recordings from MPO^Nts;VGAT-/-;hChR2^ neurons confirmed the activation of an inward current in postsynaptic neurons in response to photostimulation.

In terms of local neuronal networks, MPO^Nts^ neurons did not present reciprocal connections instead they appeared to synapse onto MPO^PACAP^ neurons. PACAP has been identified as a marker of preoptic thermoregulatory neurons activated in a warm environment (9) and also during torpor (28). Activation of MPO^PACAP^ neurons induces hypothermia (9, 28). However, PACAP is expressed in several subpopulations of neurons, representing a large proportion of the entire preoptic population (7, 28). Recent studies have identified a subpopulation of the MPO^PACAP^ neurons that expresses the EP3 prostanoid receptors, is activated in warm environment and inhibited in a cold environment (8). This population plays a critical role in the initiation of the fever response (6, 8). Modulating the activity of preoptic EP3 neurons up or down induces hypothermia or hyperthermia, respectively (8). It is possible that MPO^Nts^ neurons overlap with a previously hypothesized population of inhibitory neurons that receive peripheral cold input and project to EP3 neurons to trigger thermogenesis and inhibit heat loss(1, 2).

In view of the potent hypothermic properties of Nts when applied peripherally, centrally or in the preoptic area it was surprising to find that photostimulation of MPO^Nts;hChR2^ neurons *in vivo* resulted in a mild hyperthermia, instead. In contrast, photostimulation of MPO^Nts;VGAT-/-;hChR2^ neurons, which caused a net excitation of MPO^PACAP^ neurons, resulted in hypothermia, supporting the idea that their excitation induces hypothermia (9, 28).

Numerous studies have revealed that, in the preoptic area, GABA exerts complex effects on CBT depending on the environmental context and the specific neuronal subpopulation involved (31–34). Nts^VGAT-/-^ mice displayed significantly different thermoregulatory characteristics suggesting that GABA release from neurotensinergic neurons plays an important role in the respective responses. The fever response to endotoxin, when compared with the control, was characterized by lower CBT during the intermediate and late phases of the fever suggesting that GABA signaling is involved in the hyperthermic mechanisms activated. This finding is in line with the previous reports that preoptic GABA can induce thermogenesis (32, 33) however it is in disagreement with observations that the induction of fever is associated with decreased synaptic inhibition in the preoptic area (35, 36). It is possible that both mechanisms influence CBT during fever with the former playing a prevalent role in the intermediate and later phases.

In a warm environment preoptic release of GABA decreases and exogenous GABA-A antagonists applied locally have hyperthermic effects (32), thus Nts^VGAT-/-^ would be expected to have a higher CBT during a heat challenge. Indeed, we observed that during heat exposure the CBT of Nts^VGAT-/-^ mice increased significantly faster than in the controls suggesting that in a warm environment GABA release from neurotensinergic neurons is involved in the control of heat loss mechanisms.

During cold exposure Nts^VGAT-/-^ mice displayed higher CBT relative to controls. Previous studies have revealed increased endogenous release of GABA in the preoptic area during cold exposure and hypothermic effects of exogenous GABA-A antagonist in a cold environment (32). Thus, a decreased release of GABA from Nts^VGAT-/-^ neurons would be expected to induce a larger hypothermia during cold exposure, the opposite of the observed results. However, while an increase in preoptic GABA concentration induces hyperthermia, at all ambient temperatures, a widespread central increase in GABA is hypothermic (31, 37), therefore it is possible that VGAT deletion in Nts neurons outside of the MPO may impact CBT during cold exposure and/or that the respective regions are differentially recruited during distinct thermoregulatory mechanisms.

Finally, Nts^VGAT-/-^ mice have entered the “up” phase of the circadian CBT rhythm earlier and ended it later when compared to the control, supporting a role of neurotensinergic neurons, possibly in the suprachiasmatic nucleus, in the control of circadian rhythm (38, 39).

To question whether stimulation of all central neurotensinergic neurons induces hypothermia we have generated Nts^hM3D(Gq)^ mice. Chemogenetic activation of central Nts^hM3D(Gq)^ neurons resulted in a robust hypothermia, comparable to that induced by central infusion of exogenous neurotensin. The response was largely blocked by NtsR1 and NtsR2 antagonists injected in the MPO, suggesting that the locus of action is the MPO. Taken together our results suggest that stimulation of neurotensinergic neurons in the MPO is not sufficient to induce hypothermia although this region represents the locus where the hypothermia is triggered.

Experiments at the cellular level showed that locally applied CNO in slices from Nts^hM3D(Gq)^ mice had a net excitatory effect in nearby MPO PACAP neurons. In addition to the actions observed during photostimulation of MPO^Nts;hChR2^, namely increased sIPSCs frequency and an inward current, CNO also increased the frequency and amplitudes of sEPSCs recorded in MPO^PACAP^ neurons. This may be due to activation of glutamatergic projections from brain regions that contain glutamatergic Nts^hM3D(Gq)^ neurons. Such neurons have been identified in the posterior thalamus and in the ventrolateral periaqueductal gray (19, 20). It is also possibIe that some of the parabrachial nucleus neurons’ glutamatergic projections (40) are also neurotensinegic. It is likely that the chemogenetic stimulation of MPO Nts^hM3D(Gq)^ neurons and projections elevates the local Nts concentration to a level that activates the low affinity NtsR2. These receptors are expressed in MPO astrocytes and upon activation stimulate glutamate release from nearby synaptic terminals as observed when exogenous Nts is applied locally (24). A schematic model of the network and of the cellular mechanisms involved in the CBT modulation by neurotensinergic neurons is presented in Fig 8.

**Figure 8.**
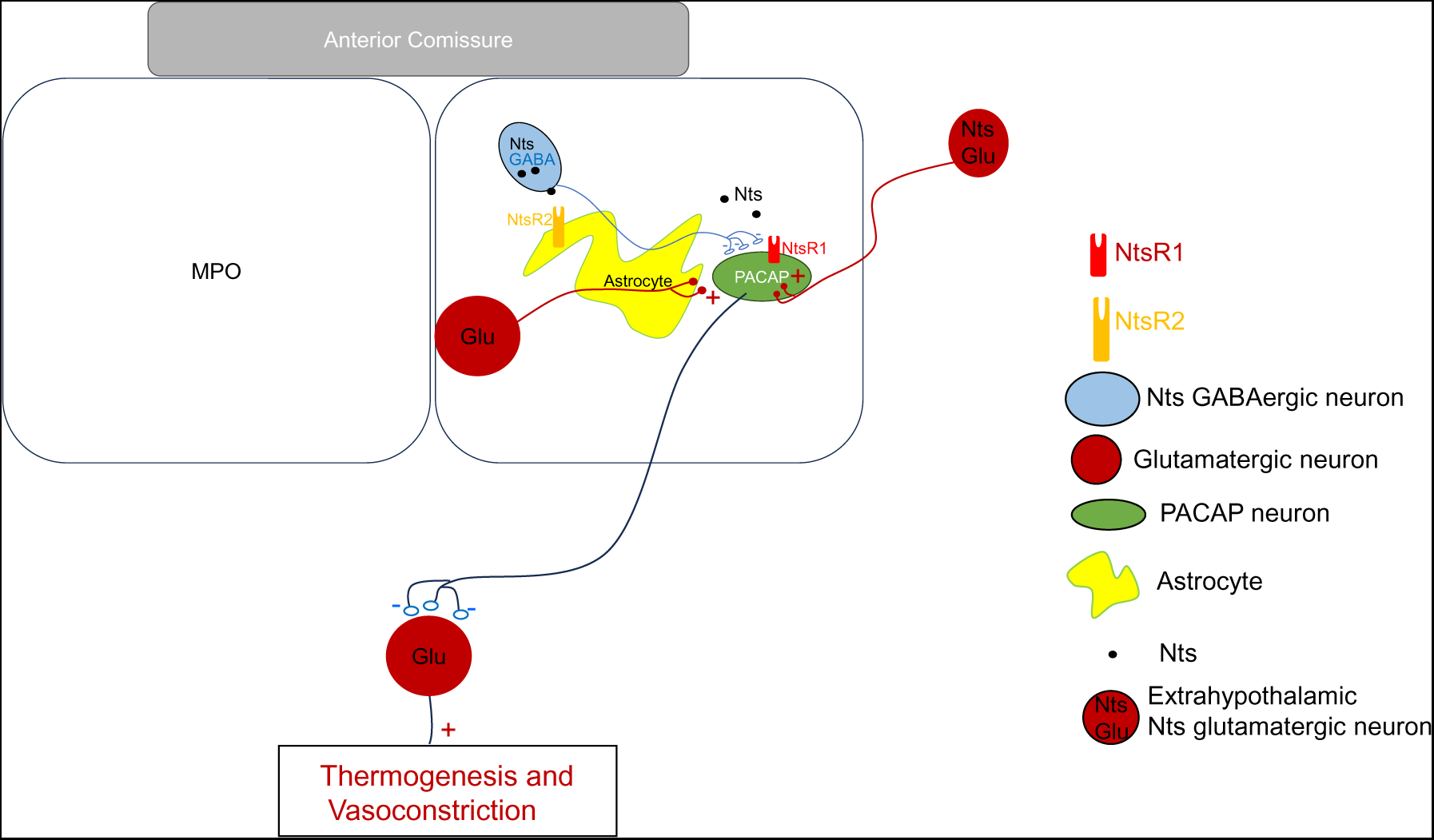
Schematic representation of neurotensinergic neurons in a thermoregulatory preoptic pathway. Preoptic thermoregulatory PACAP neurons, assumed to be glutamatergic, project to inhibitory interneurons in other brain regions that project to neurons controlling thermogenesis and vasoconstriction. PACAP neurons’ inhibition results in increased thermogenesis and decreased vasoconstriction resulting in increased CBT. Conversely, excitation of the PACAP thermoregulatory neurons results in hypothermia. Preoptic Nts neurons are GABAergic and project to preoptic thermoregulatory PACAP neurons and modulate their activity. Preoptic astrocytes express NtsR2 receptors and their activation modulates the release of glutamate from nearby synaptic terminals.

In summary, this study characterizes the cellular properties of MPO^Nts^ neurons and their influence on CBT and provides insights in the cellular mechanisms at the MPO level that result in hypothermia.

## Materials and Methods

### Animals

Experiments on animals were carried out in accordance with the National Institute of Health Guide for the care and use of Laboratory animals (1996 (7th ed.) Washington DC: National Research Council, National Academies Press). The protocols were reviewed and approved by the Institutional Animal Care and Use Committee of the Scintillon Institute. The standards are set forth by the American Association for the Accreditation of Laboratory Animal Care (AAALAC) and in the Animal Welfare Act. The work was designed in such a way to minimize the number of animals used as well as their suffering.

The Nts-cre driver line (B6;129-Ntstm1(cre)Mgmj/J; stock no: 017525) (23) was purchased from Jackson Laboratory (Bar Harbor, ME, USA). The Nts-cre driver line was crossed with the Slc32a1tm1Lowl (also referred to as VGATflox/flox) (Jackson Laboratory; stock no: 012897) to generate mice that have the GABA vesicular transporter VGAT (solute carrier family 32 member 1), knocked-down in Nts neurons. Heterozygous double transgenic mice Nts+/- ;Vgat+/-were crossed to obtain Nts+/+;Vgat-/-. These mice are referred to as Nts^VGAT-/-^. We have also crossed the Nts-cre driver line with the B6N;129-Tg(CAG-CHRM3*,-mCitrine)1Ute/J line (Jackson Laboratory; stock no: 026220) to generate mice that express the excitatory designer receptor hM3D(Gq) in Nts-neurons in all brain regions. This new line was named Nts^hM3D(Gq)^. In chemogenetics experiments, to activate hM3D(Gq), mice received an i.p. injection of CNO (0.7-0.8 mg/kg).

### Transgenic animals and AAV injections

The Nts-cre homozygous mice (4 to 6 months old) received bilateral stereotaxic injections (200 nL at a rate of 0.1 μL/min) of AAV-EF1a-double floxed-hChR2(H134R)-eYFP-WPRE-HGHpA (Addgene #20298; 2.1 x 10^13^ virus molecules/ml) or AAV5-EF1a-DIO-eYFP (Addgene #27056; 1.3 x 10^13^ virus molecules/ml) in the MPO to express the opsin Channelrhodopsin2 (hChR2) or eYFP (control) in MPO Nts neurons. We refer to these mice as MPO^Nts;hChR2^ and MPO^Nts;eYFP^, respectively. To ensure that only cre-expressing neurons were targeted we have carried out control injections of the two viral vectors in wild-type C57/Bl6 mice (3 mice for each) and confirmed that no fluorescent signal was detected in the brain of these mice. We have carried out RNAscope for Nts (see below) to determine the colocalization with eYFP (Fig 9) in 3 MPO^Nts;hChR2^ mice. We have studied colocalization from 6 randomly chosen field of view from 3 slices from 3 different animals and found colocalization in 82% (355 out of 433) of the Nts positive neuros. Only 4 out of 359 eYFP neurons appeared to be Nts-negative (1.1%). Together with the detection of Nts transcripts in the labeled neurons (see Results section) we conclude that we have specifically targeted Nts neurons using these viral vectors.

**Figure 9.**
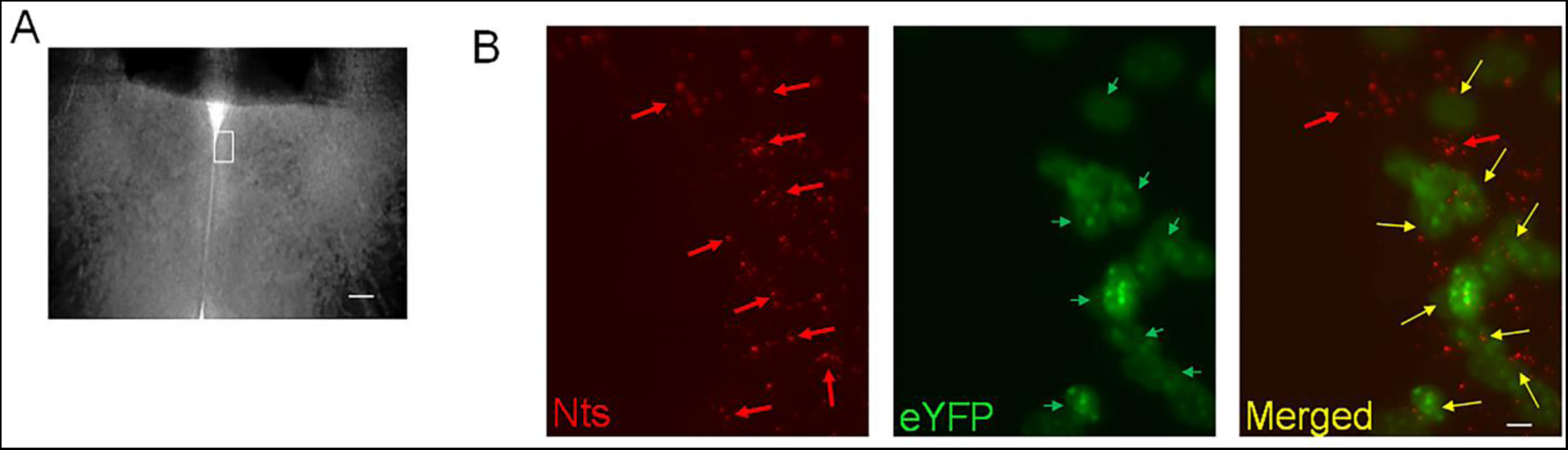
Transduction of MPO^Nts^ neurons with ChR2-eYFP by injecting AAV-EF1a-double floxed-hChR2(H134R)-eYFP in Nts-cre mice. **A.** Brightfield image of a MPO slice. The white rectangle represents the region imaged in **B**. The scale bar represents 100 µm. **B.** Representative images of Nts transcripts visualized using RNAscope (red, left panel), eYFP (green, middle) and their superimposed images (right) in a coronal slice from a MPO^Nts;hChR2^ mouse. eYFP was visible in 8 out of 10 Nts positive cells. The scale bar represents 10 µm.

Finally, we have performed bilateral stereotaxic injections (200 nL at a rate of 0.1 μL/min) of the AAV5-EF1a-double floxed-hChR2(H134R)-eYFP-WPRE-HGHpA (Addgene #20298; 2.1 x 10^13^ virus molecules/ml) or AAV5-EF1a-DIO-eYFP (Addgene #27056; 1.3 x 10^13^ virus molecules/ml) in the MPO of Nts^VGAT-/-^ to express ChR2 or eYFP, in MPO Nts^VGAT-/-^ neurons. We refer to these mice as MPO^Nts;VGAT-/-;hChR2^ and MPO^Nts;VGAT-/-;eYFP^, respectively.

All experiments were carried out in male mice. Key sets of experiments were also performed in female mice as specified in the text.

### Estrous cycle monitoring

Female mice have a 4–6 day estrous cycle that consists of four stages – proestrus, estrus, diestrus I/metestrus, and diestrus/II. Mice were habituated to handling and vaginal smears prior to the behavioral experiments. Mice were swabbed daily before 9a.m. and experiments started after 10 am. The stage of the cycle was determined using vaginal cytology. The smears were stained with hemalum-eosin and analyzed microscopically. Experiments in females were carried out in diestrus days since during this stage the baseline CBT is less variable for each animal and among different animals. The days of diestrus are characterized by the presence of leukocytes and the absence of keratinized epithelial cells with no nuclei.

### Telemetry and MPO Injections

For CBT measurements the mice (4 to 6 months old) were anesthetized with isoflurane (induction 3%-5%, maintenance 1%-1.5%) and radio telemetry devices (Anipill, BodyCap, Hérouville Saint-Clair, France) were surgically implanted into the peritoneal cavity. For MPO injections, mice were stereotaxically implanted with a bilateral guide cannula (27 Ga) as described in our previous studies (41). Coordinates for cannula implants in the MPO were: from Bregma: 0.05 mm, 0.35 mm, and −0.35 mm lateral, and ventral 4.75 mm (42).The ambient temperature was maintained at ∼28 ± 0.5°C in a 12:12-hour light-dark cycle-controlled room (lights on 8:00 am, ZT0). All substances injected were dissolved in sterile artificial cerebrospinal fluid (aCSF). For MPO injections, mice were placed in a stereotaxic frame and the injector (33 Ga) was lowered inside the cannula. A volume of 100 nL (rate 0.1 μL/min) was delivered using an injector connected to a microsyringe (0.25 μL). After injections the animal was returned to the home cage. Injections were carried out at 10 am local time, during the “subjective light period.”

### Slice Preparation

Coronal tissue slices containing the MPO were prepared from mice 4-6 months old MPO^Nts;hChR2^, MPO^Nts;VGAT-/-;hChR2^ or Nts^hM3D(Gq)^ mice. The slice preparation was as previously described (41). The slice used in our recordings corresponded to the sections located from 0.15 mm to -0.05 mm from Bregma in the mouse brain atlas (42).

### Whole-cell patch-clamp recording

Whole-cell patch-clamp clamp was performed as described in our previous studies (41). The aCSF contained (in mM) the following: 130 NaCl, 3.5 KCl, 1.25 NaH_2_PO_4_, 24 NaHCO_3_, 2 CaCl_2_, 1 MgSO_4_, and 10 glucose, osmolarity of 300–305 mOsm, equilibrated with 95% O_2_ and 5% CO_2_, pH 7.4. Other salts and agents were added to this saline. Whole-cell recordings were carried out using a K^+^ pipette solution containing (in mM) 130 K-gluconate, 5 KCl, 10 HEPES, 2 MgCl_2_, 0.5 EGTA, 2 ATP and 1 GTP (pH 7.3). The electrode resistance after back-filling was 2–4 MΩ. All voltages were corrected for the liquid junction potential (−13 mV). Data were acquired with a MultiClamp 700B amplifier (Molecular Devices, Sunnyvale, CA, USA) digitized using a Digidata 1550 interface and the Pclamp10.6 software package. The sampling rate for the continuous recordings of spontaneous activity was 50 kHz. The cell capacitance was determined and compensated using the Multiclamp Commander software.

The recording chamber was constantly perfused with extracellular solution (2–3 mL·min^−1^). The antagonists were bath-applied. The bath temperature was maintained at 36–37°C by using an inline heater and a TC-344B temperature controller (Warner Instruments, Hamden, CT, USA).

Synaptic activity was quantified and analyzed statistically as described previously (41, 43). Synaptic events were detected and quantified (amplitude, kinetics, frequency) off-line using a peak detection program (Mini Analysis program, Synaptosoft, Decatur, NJ, USA). Events were detected from randomly selected recording stretches of 2 min before and during incubation with pharmacological agent. Cumulative distributions of the measured parameters (inter-event interval, amplitude, rise time, time constant of decay) were compared statistically using with the Kolmogorov-Smirnov two-sample test (K-S test, P<0.05) using the Mini Analysis program. The averages for the measured parameters (frequency, amplitude, rise time, time constant of decay) for each experiment were obtained using the Mini Analysis program. Event frequency was calculated by dividing the number of events by the duration (in seconds) of the analyzed recording stretch.

Spot illumination of MPO^Nts;hChR2^ neurons was carried out using a Polygon300 illumination system (Mightex, Toronto, Canada) that allows the control of the size, shape, intensity and position of the illuminated spot as described previously. In a field of view there were 1-4 fluorescent neurons. The position and size of the spot were controlled using the PolyScan2 software (Mightex). The intensity of the light at the bottom of the recording chamber was measured using a photosensor (ThorLabs, Newton, NJ, USA) and ranged from 1.2 to 1.4 mW/mm^-2^. The light pulses were 50 ms long and delivered at 10Hz for durations of 20 to 120 s.

### Optogenetic stimulation *in vivo*

A dual fiber-optic cannula (100µm diameter, NA 0.22, 0.7 mm pitch,) (Doric Lens, Quebec, Canada) was implanted as described above for the guide cannula. Light pulses were applied using high-power photodiodes, a digital and analog I/O control module and Polygon 300 software (Mightex, Toronto, Canada) via dual fiber cables (Doric Lens, Quebec, Canada). The intensity of the light at the end of the optic cannula was measured using a fiber photodiode power sensor (ThorLabs, Newton, NJ, USA) and ranged from 3.9 to 4.3 mW/mm^-2^. The light pulses were 50 ms long and delivered at 10Hz for 30 s periods followed by 10s recovery breaks. The total duration of light stimulation was 60 min.

### Chemicals

Agonists and antagonists were purchased from Tocris (Ellisville, MO, USA). The other chemicals were from Sigma (Carlsbad, CA, USA).

### Cell Harvesting, Reverse Transcription and PCR

MPO neurons in slices were patch-clamped and then harvested into the patch pipette by applying negative pressure as previously described (41, 43). The content of the pipette was expelled in a PCR tube. dNTPs (0.5 mM), 50 ng random primers (Invitrogen) and H_2_O were added to each cell to a volume of 16 μl. The samples were incubated at 65°C for 5 min and then put on ice for 3 min. First strand buffer (Invitrogen), DTT (5 mM, Invitrogen), RNaseOUT (40 U, Invitrogen) and SuperScriptIII (200 U, Invitrogen) were added to each sample to a volume of 20 μl followed by incubation at room temperature for 5 min, at 50°C for 50 min and then at 75°C for 15 min. After reverse transcription samples were immediately put on ice. 1 μl of RNAse H was added to samples and kept at 37°C for 20 min. PCR assays were carried out using the pairs of primers listed in Suppl Table 1. The PCR products were verified using Sanger sequencing. Two rounds of PCR, using nested primers, were used to detect Nts and Vglut2 transcripts. For PACAP and VGAT a second round of PCR was not necessary since it did not yield additional positives when compared with the first round. Two negative controls were routinely carried out. The first one was amplified from a harvested cell without reverse-transcription while the second was represented by a RT/PCR of the “pipette tip” (e.g. obtained when a successful gigaseal was not achieved and the cytoplasm was not harvested by suction). Since all single MPO cells tested were negative to Vglut2 a second positive control was carried out. Ventromedial hypothalamic neurons, a predominantly glutamatergic population was tested. The positivity rate was 90% (9 of 10 neurons tested) (Suppl Fig1B).

### RNAscope assays

Tissue was processed using in situ hybridization to detect mRNA for Nts, VGAT (Slc32a1). 40-μm thick coronal sections fixed in 4% PFA for 15 min at 4°C, dehydrated in serial concentrations of ethanol (50%–100%), and processed according to the protocol provided in the RNAscope kit (ACDbio Cat# 320293). Sections were hybridized with the following mixed probes; Nts (Mm-Nts, Cat. 420441) and VGAT (Mm-Slc32a1-C2, Cat. 319191-C2) for 2 h at 40°C.

### Endotoxin induced fever, heat and cold exposure tests

To induce a fever response we injected i.p. a dose of 0.1 mg/kg lipopolysaccharides (LPS) dissolved in 0.3 ml sterile saline. To study the CBT change to heat exposure the mice were placed, in their home cages, in a 37 °C incubator. Their CBT was monitored continuously by telemetry, and an animal was removed from the incubator once it reached a CBT of 41.5 °C. For cold exposure tests the mice were placed in a cold room (4 °C) for 3h. All the experiments were started at 10 a.m. (ZT2).

### Data quantification and statistical analysis

The values reported are presented as mean ± standard deviation (SD). We used power analyses (www.biomath.info) with values from our data (means and SDs) to calculate that our study was powered to detect a 0.7 °C change in CBT with at least >80% reliability for all transgenic models used in this study. The normality of the samples was tested using the Shapiro-Wilk and Anderson-Darling tests. Statistical significance of the results pooled from two groups was assessed with t-tests or Mann-Whitney test using Prism9 (GraphPad Software). One-way analysis of variance (ANOVA, Kruskal-Wallis) with Tukey’s post hoc test (P<0.05) was used for comparison of multiple groups. Cumulative distributions were compared with the Kolmogorov-Smirnov test (P<0.05). Data collected as time series were compared across time points by one-way ANOVA with repeated measures (P<0.05) (Prism4, GraphPad Software), followed by Mann Whitney U tests (P<0.05) for comparisons at each time point. The statistical value, the degrees of freedom and the p value are reported in the figure legends or, if data is not presented in a figure, the respective values are reported in the text. P values for the results of Tukey’s tests are presented in tables.

### Key resources table

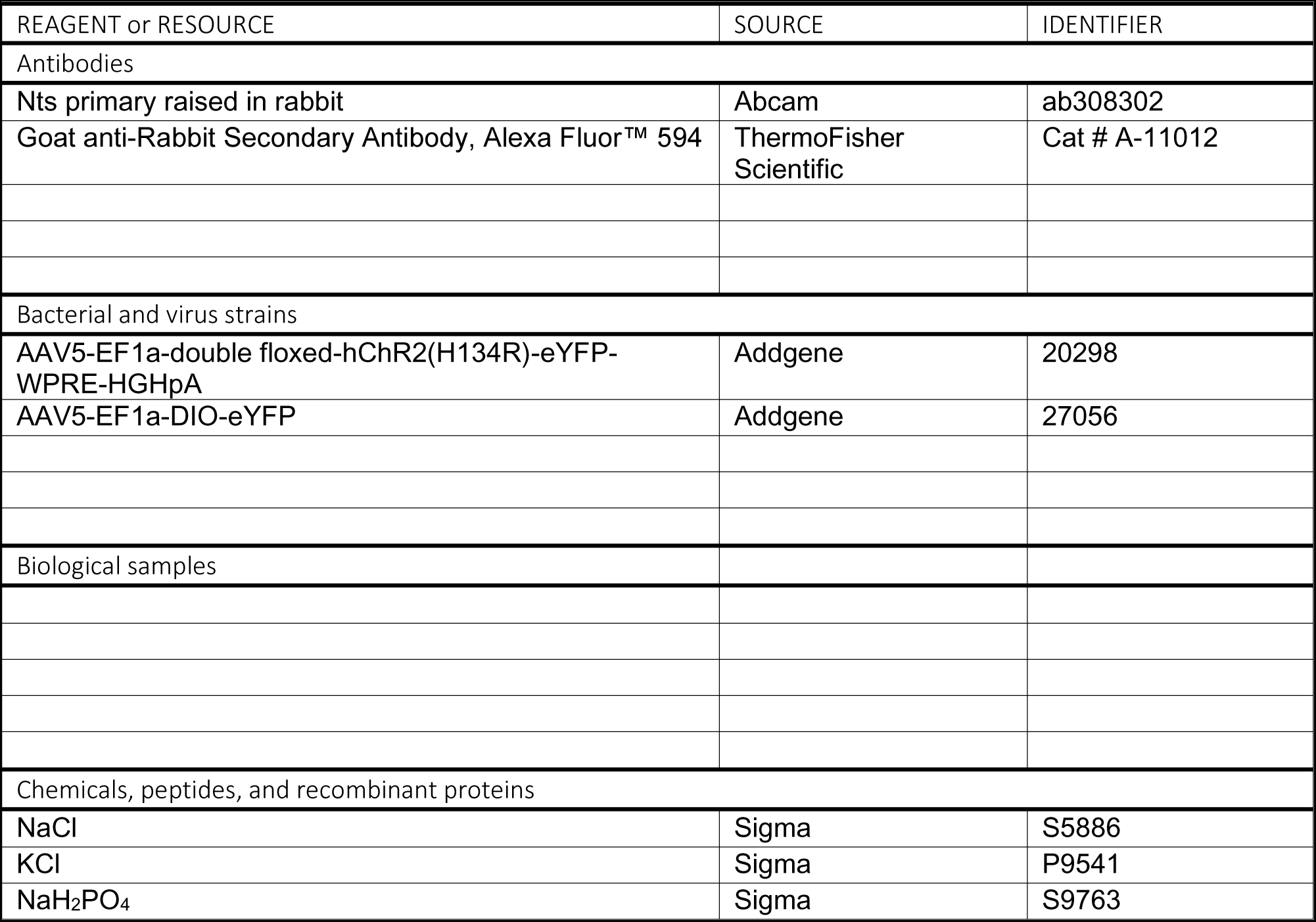
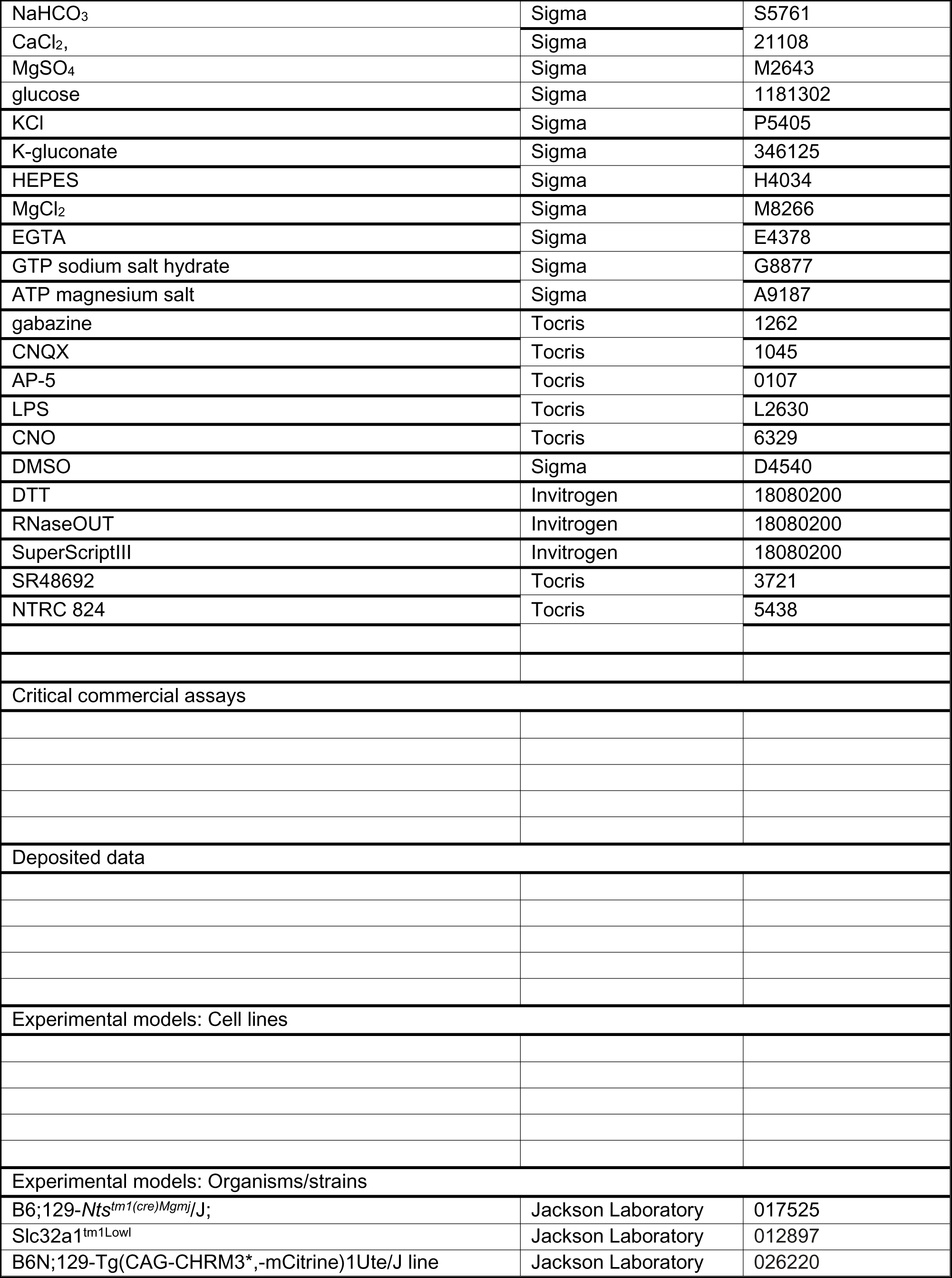
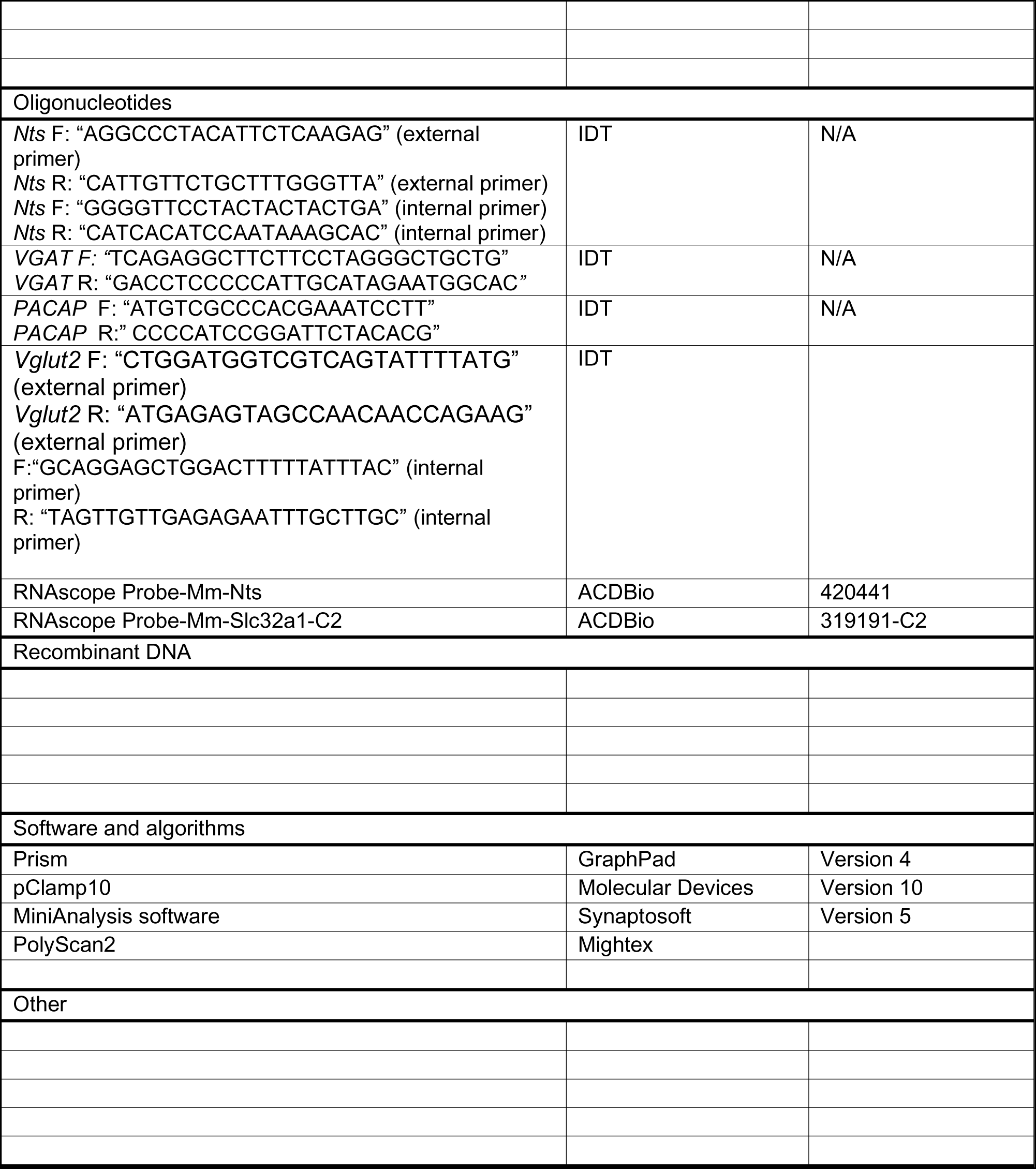

## Acknowledgments

This is manuscript number 1073 from the The Scintillon Institute. This research was supported by the National Institutes of Health Grants NS094800 and NS124844 (I.V.T.).

---

**Figure 1 Suppl.**
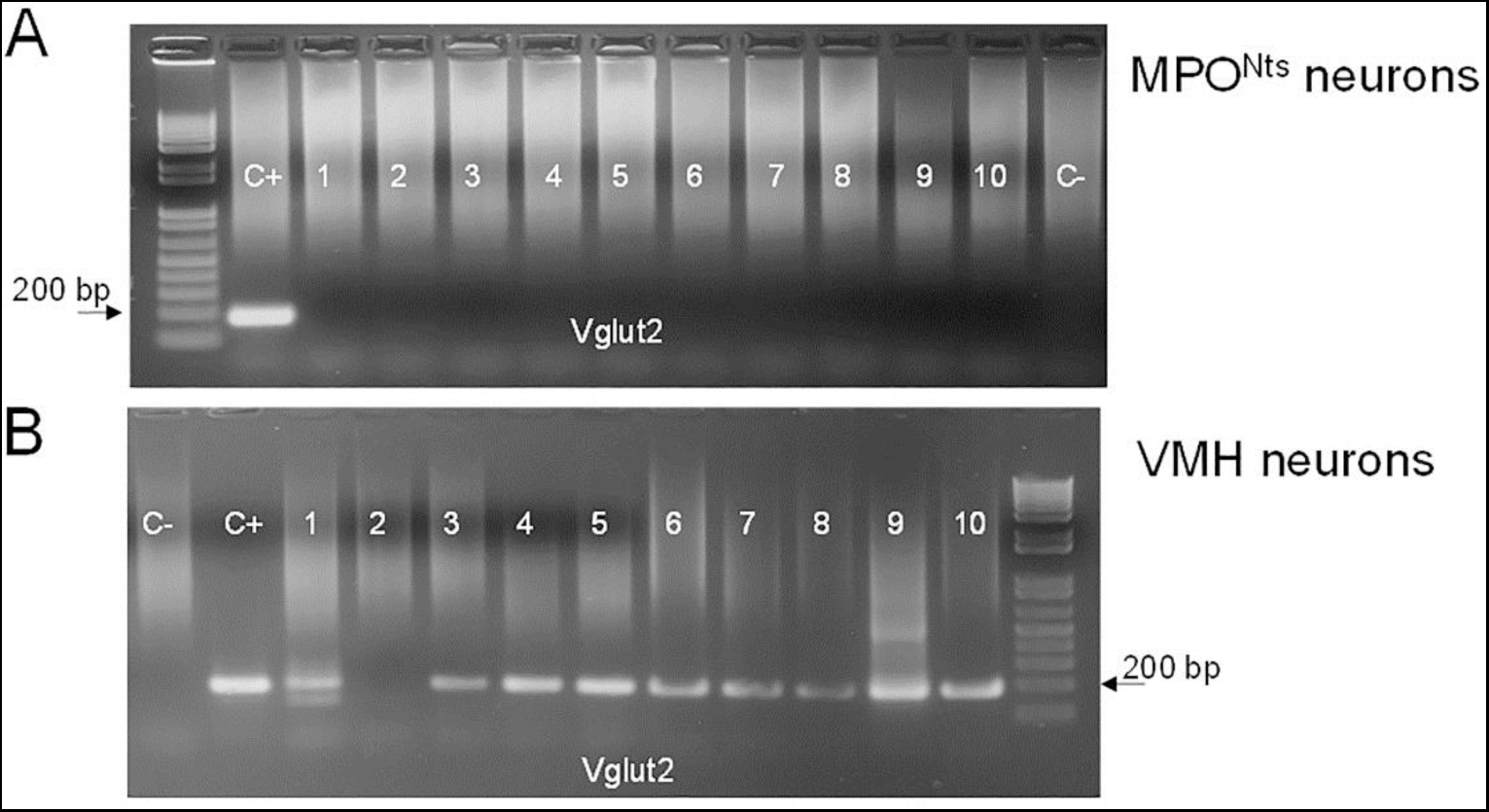
Lack of Vglut2 expression in MPO^Nts^ neurons. **A.** Single cell RT/PCR analysis of Vglut2 expression. Representative results from 10 MPO^Nts;hChR2^ neurons. **B.** Vglut2 expression in single ventromedial hypothalamus (VMH) neurons. Vglut2 transcripts were detected in 9 out of 10 neurons. **A,B.** The expected size of the PCR product is 184 base pairs. Negative (−) control was amplified from a harvested cell without reverse-transcription, and positive control (+) was amplified using 1 ng of hypothalamic mRNA.

**Figure 3 Suppl.**
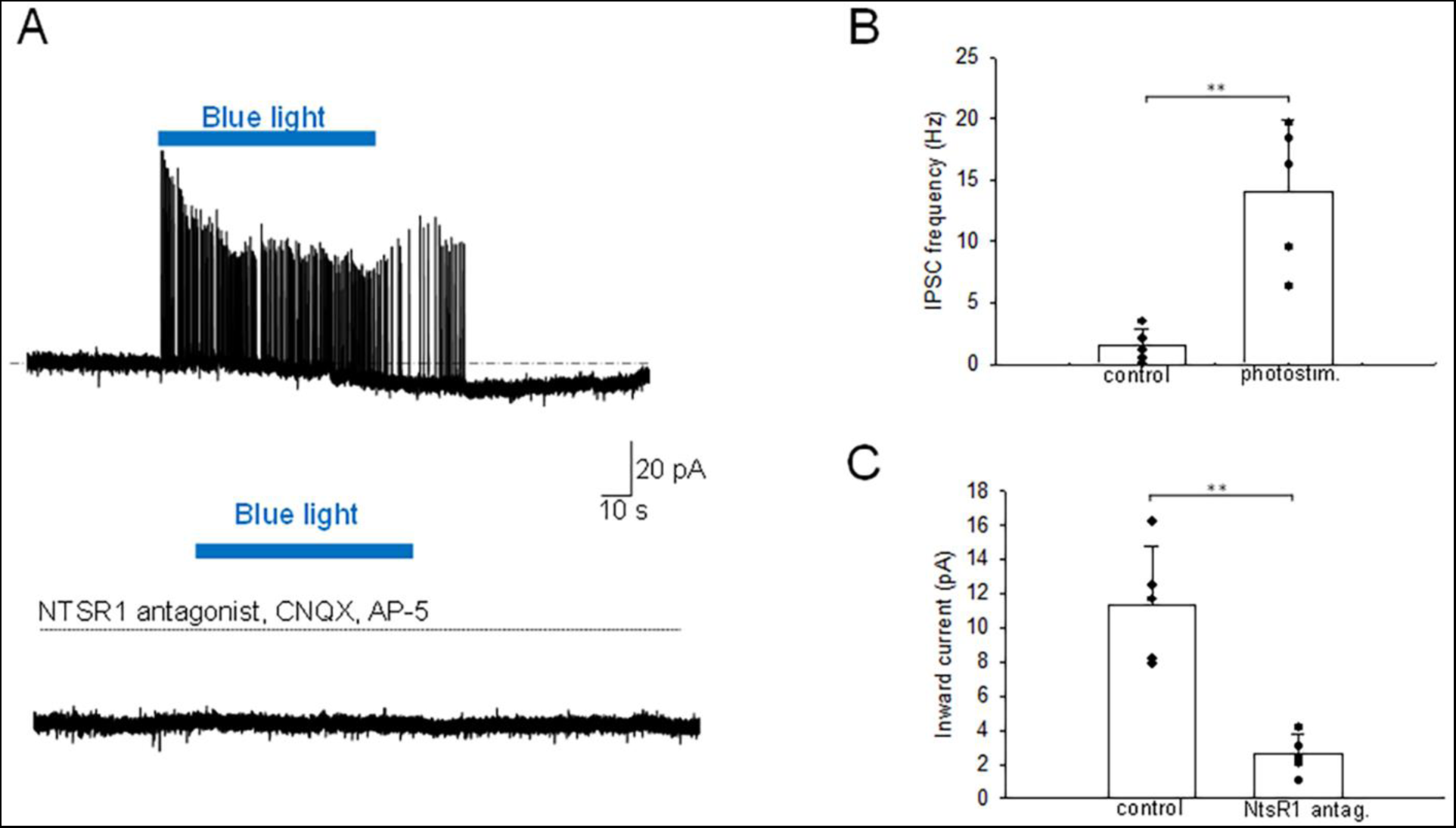
Optogenetic stimulation of MPO^Nts;hChR2^ neurons from females increases the frequency of IPSCs and activates an inward current in nearby MPO neurons. **A.** Optogenetic stimulation of a MPO^Nts;hChR2^ neuron activates IPSCs and an inward current in a nearby MPO neuron (upper trace). In the presence of CNQX (20 µM), AP-5 (50 µM), Gabazine (5 µM) and the NtsR1 antagonist SR48692 (100 nM) light stimulation was without effect (lower trace). The neuron was held at -50 mV. **B,C.** Bar charts summarizing the increase in the frequency of IPSCs (**B**) and the amplitude of the inward current (**C**) recorded in MPO neurons in response to optogenetic stimulation of several MPO^Nts;hChR2^ neurons. **B.** The IPSCs frequency increased from1.5±1.4 Hz to 14.1±5.8 Hz in response to photostimulation (one-way ANOVA F(1,9)=22.4, p=1.4x10^-3^). **C**. The average inward current activated by optogenetic stimulation decreased from 11.3±3.4 pA to 2.6±1.2 pA in the presence of the NtsR1 antagonist SR48692 (100 nM) (one-way ANOVA (F(1,9)=28.9, p=6.6x10^-4^). Bars represent means ± S.D. Data pooled from n=5 neurons.

**Figure 4 Suppl.**
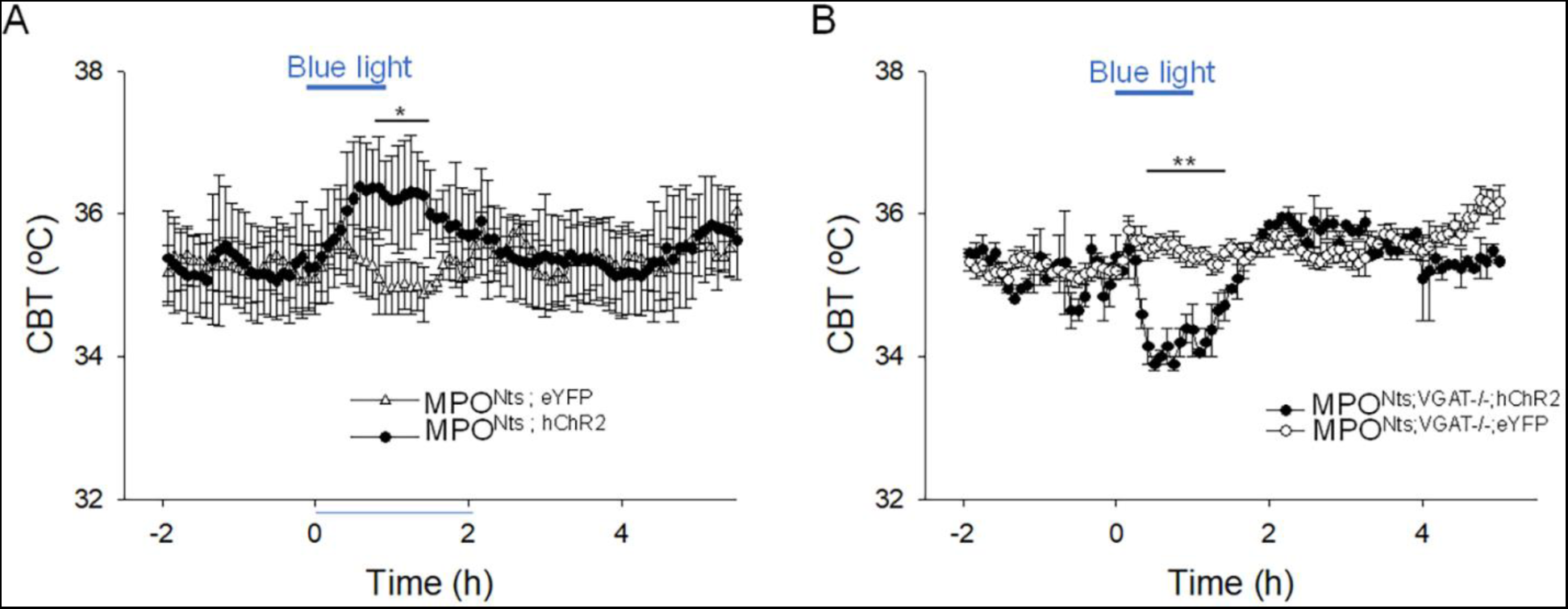
In female mice optogenetic activation of MPO^Nts;hChR2^ neurons induces hyperthermia while optogenetic activation of MPO^Nts;VGAT-/-;hChR2^ neurons induces hypothermia. **A.** Optogenetic stimulation of MPO^Nts;hChR2^ neurons (●) *in vivo* for 1 hour (blue light) induced a hyperthermia of 1.31±0.65 °C relative to control (Δ). The response was statistically different to the response to photostimulation of control MPO^Nts;eYFP^ mice (Δ) (one-way repeated measures ANOVA, F(1,110)=5.7, p=1.9x10^-2^, followed by Man-Whitney U tests for each time point, * P<0.05). **B.** Optogenetic stimulation of MPO^Nts;VGAT-/-;hChR2^ neurons (●) *in vivo* for 1 hour (blue light) induced a hypothermia of 1.69±0.22 °C relative to control (Δ). The response was statistically different to the response to photostimulation of control MPO^Nts;VGAT-/-;eYFP^ mice (Δ) (one-way repeated measures ANOVA, F(1,83)=9.9, p=2.2x10^-3^, followed by Man-Whitney U tests for each time point, ** P<0.01). **A,B.** The points represent averages±S.D. through the 7h recording period. Experiments were carried out in parallel in groups of 6 female mice.

**Figure 5 Suppl.**
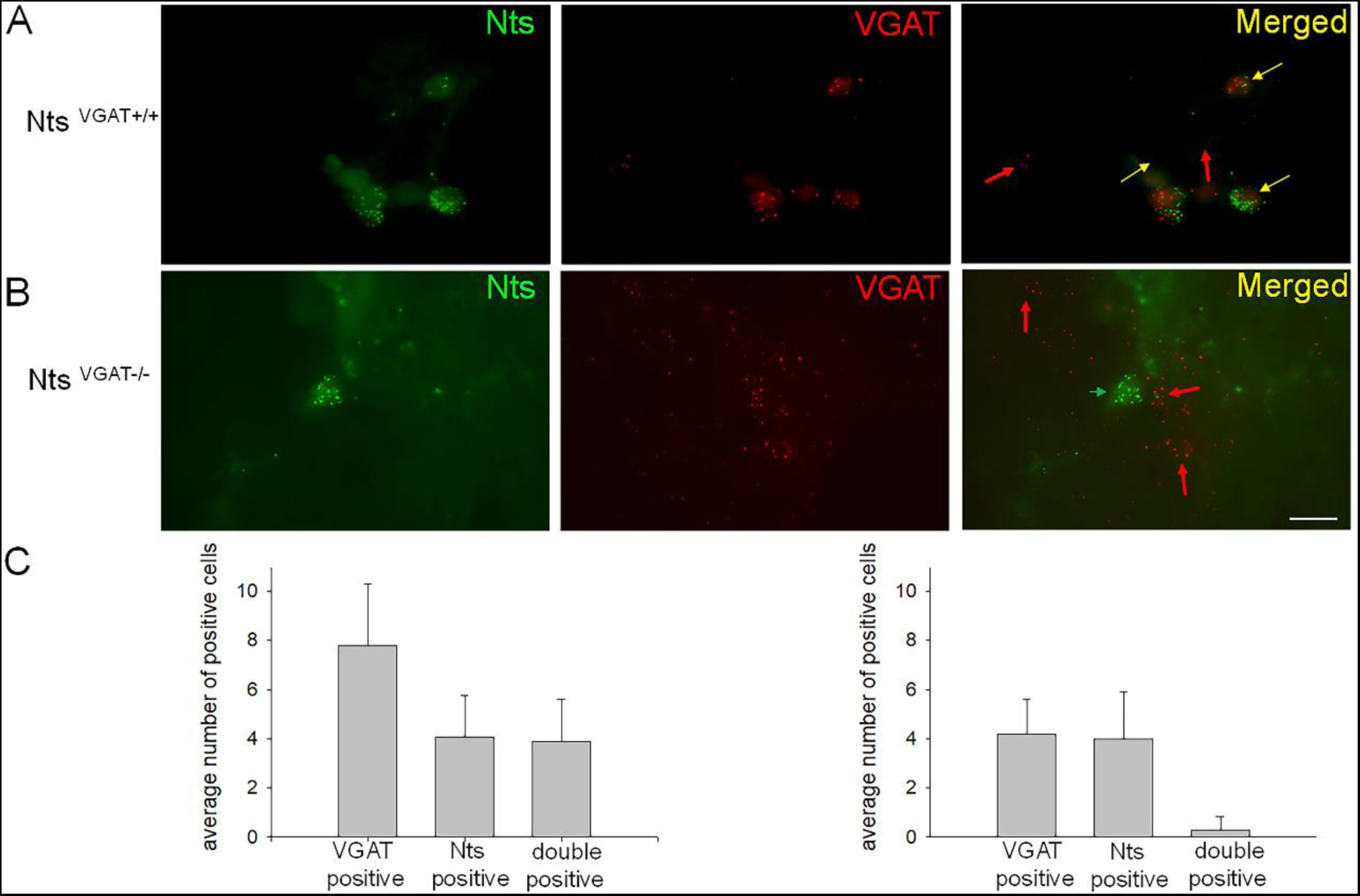
Expression of VGAT transcripts in preoptic slices from Nts^VGAT-/-^ and Nts^VGAT+/+^ male mice. **A,B.** Representative images of VGAT (red) and Nts (green) transcripts visualized using RNAscope technology in preoptic slices from Nts^VGAT+/+^ (**A**) and Nts^VGAT-/-^ mice (**B**). **A.** Nts transcripts (green, left) are present in 3 out of 5 VGAT positive cells (red, middle) as indicated by their superimposed images (right, yellow arrows). Two other VGAT expressing cells do not express Nts transcripts (right, red arrows). **B.** Nts transcripts (green, left) are present in 2 cells while VGAT transcripts (red, middle) are present in 6 cells (middle). In Nts^VGAT-/-^ tissue the VGAT positive cells (right, red arrows) do not co-express Nts transcripts. **A,B.** The scale bar represents 10 µm. **C.** Bar charts summarizing the average number positive cells for the respective transcripts in a randomly selected field of view in Nts^VGAT+/+^ (left) and Nts^VGAT-/-^ (right) preoptic slices. Data were averaged from 15 randomly selected fields of view and 3 different mice for each genotype. Overall, 139 cells expressed VGAT, 72 cells expressed Nts transcripts and 68 cells expressed both in Nts^VGAT+/+^ tissue. In the Nts^VGAT-/-^ tissue 75 cells expressed VGAT, 72 cells expressed Nts transcripts and 3 cells expressed both transcripts. The percentage of Nts expressing cells that co-expressed VGAT was 94.4% and 4.2% in Nts^VGAT+/+^ and Nts^VGAT-/-^ tissue, respectively.

**Supplementary Table 1.**
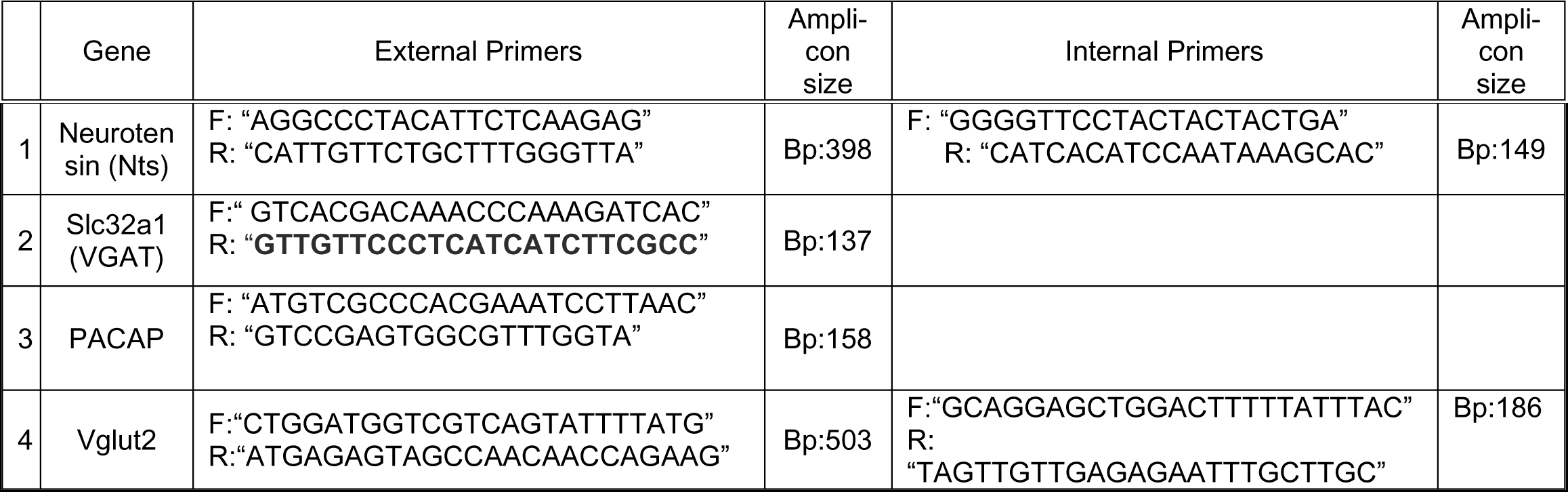
PCR primers for the genes studied.

**Supplementary Table 2.** P-values for Tukey’s test comparisons among groups (Fig 2B)

| Pair | P-value |
| --- | --- |
| X<sub>1</sub>-X<sub>2</sub> | 0.0001476 |
| X<sub>1</sub>-X<sub>3</sub> | 0.0003445 |
| X<sub>1</sub>-X<sub>4</sub> | 0.0005521 |
| X<sub>1</sub>-X<sub>5</sub> | 0.007513 |
| X<sub>1</sub>-X<sub>6</sub> | 0.01494 |
| X<sub>2</sub>-X<sub>3</sub> | 1.676e-8 |
| X<sub>2</sub>-X<sub>4</sub> | 1.751e-7 |
| X<sub>2</sub>-X<sub>5</sub> | 1.178e-8 |
| X<sub>2</sub>-X<sub>6</sub> | 1.447e-7 |
| X<sub>3</sub>-X<sub>4</sub> | 0.006143 |
| X<sub>3</sub>-X<sub>5</sub> | 0.0007085 |
| X<sub>3</sub>-X<sub>6</sub> | 0.002418 |
| X<sub>4</sub>-X<sub>5</sub> | 0.0003605 |
| X<sub>4</sub>-X<sub>6</sub> | 0.000312 |
| X<sub>5</sub>-X<sub>6</sub> | 0.2656 |

**Supplementary Table 3.** P values of the Tukey’s test comparisons among groups (Fig 5B)

| Columns | P-value |
| --- | --- |
| x1-x2 | $7.1 \times 10^{-5}$ |
| x1-x3 | 0.7237 |
| x2-x3 | $7.82 \times 10^{-6}$ |

**Supplementary Table 4.** P values of the Tukey’s test comparisons among groups (Fig 7F)

| Pair | P-value |
| --- | --- |
| x1-x2 | 0.044 |
| x1-x3 | 0.4212 |
| x1-x4 | 0.0201 |
| x2-x3 | 0.543 |
| x2-x4 | 0.978 |
| x3-x4 | 0.3303 |

**Supplementary Table 5.** P values of the Tukey’s test comparisons among groups (Fig 7G)

| Pair | P-value |
| --- | --- |
| x1-x2 | 0.0001877 |
| x1-x3 | 0.05677 |
| x1-x4 | 0.001055 |
| x2-x3 | 0.04938 |
| x2-x4 | 0.8145 |
| x3-x4 | 0.2338 |

**Supplementary Table 6.**
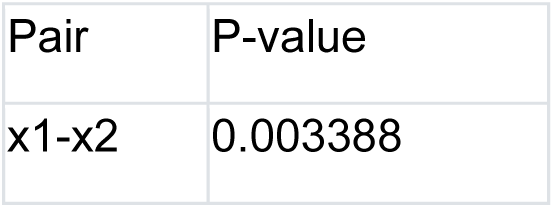
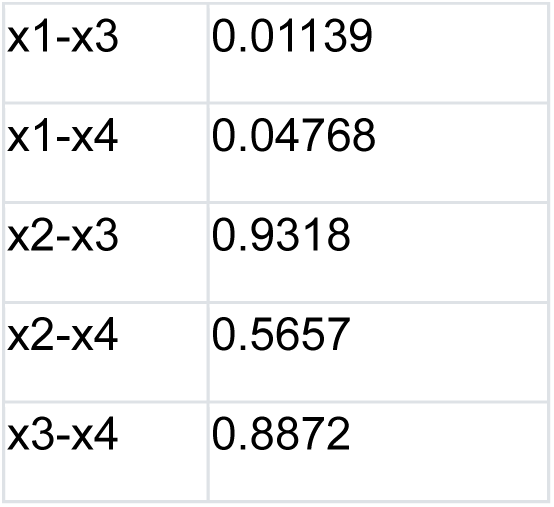
P values of the Tukey’s test comparisons among groups (Fig 7H)

**Supplementary Table 7.**
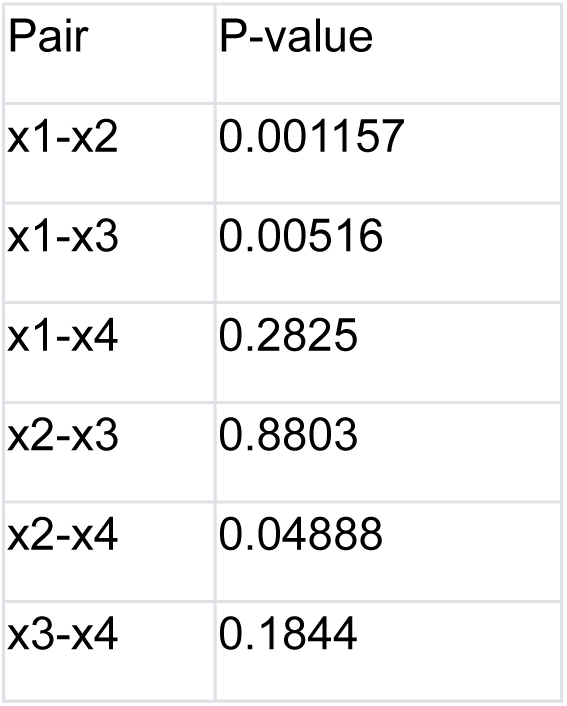
P values of the Tukey’s test comparisons among groups (Fig 7I)

| Pair | P-value |
| --- | --- |
| x1-x2 | 0.001157 |
| x1-x3 | 0.00516 |
| x1-x4 | 0.2825 |
| x2-x3 | 0.8803 |
| x2-x4 | 0.04888 |
| x3-x4 | 0.1844 |

## References

1. Morrison SF, Nakamura K. Central Mechanisms for Thermoregulation. Annu Rev Physiol. 2019;81:285–308.

2. Nakamura K, Nakamura Y, Kataoka N. A hypothalamomedullary network for physiological responses to environmental stresses. Nat Rev Neurosci. 2022;23(1):35–52.

3. Tan CL, Knight ZA. Regulation of Body Temperature by the Nervous System. Neuron. 2018;98(1):31–48.

4. Morrison SF. Efferent neural pathways for the control of brown adipose tissue thermogenesis and shivering. Handb Clin Neurol. 2018;156:281–303.

5. Saper CB, Romanovsky AA, Scammell TE. Neural circuitry engaged by prostaglandins during the sickness syndrome. Nat Neurosci. 2012;15(8):1088–95.

6. Machado NLS, Bandaru SS, Abbott SBG, Saper CB. EP3R-Expressing Glutamatergic Preoptic Neurons Mediate Inflammatory Fever. J Neurosci. 2020;40(12):2573–88.

7. Moffitt JR, Bambah-Mukku D, Eichhorn SW, Vaughn E, Shekhar K, Perez JD, et al. Molecular, spatial, and functional single-cell profiling of the hypothalamic preoptic region. Science. 2018;362(6416).

8. Nakamura Y, Yahiro T, Fukushima A, Kataoka N, Hioki H, Nakamura K. Prostaglandin EP3 receptor-expressing preoptic neurons bidirectionally control body temperature via tonic GABAergic signaling. Sci Adv. 2022;8(51):eadd5463.

9. Tan CL, Cooke EK, Leib DE, Lin YC, Daly GE, Zimmerman CA, et al. Warm-Sensitive Neurons that Control Body Temperature. Cell. 2016;167(1):47–59 e15.

10. Vincent JP, Mazella J, Kitabgi P. Neurotensin and neurotensin receptors. Trends Pharmacol Sci. 1999;20(7):302–9.

11. Schroeder LE, Leinninger GM. Role of central neurotensin in regulating feeding: Implications for the development and treatment of body weight disorders. Biochim Biophys Acta Mol Basis Dis. 2018;1864(3):900–16.

12. Kleczkowska P, Lipkowski AW. Neurotensin and neurotensin receptors: characteristic, structure-activity relationship and pain modulation--a review. Eur J Pharmacol. 2013;716(1-3):54–60.

13. Kyriatzis G, Khrestchatisky M, Ferhat L, Chatzaki EA. Neurotensin and Neurotensin Receptors in Stress-related Disorders: Pathophysiology & Novel Drug Targets. Curr Neuropharmacol. 2023.

14. Ramirez-Virella J, Leinninger GM. The Role of Central Neurotensin in Regulating Feeding and Body Weight. Endocrinology. 2021;162(5).

15. Torruella-Suarez ML, McElligott ZA. Neurotensin in reward processes. Neuropharmacology. 2020;167:108005.

16. Chen Z, Chen G, Zhong J, Jiang S, Lai S, Xu H, et al. A circuit from lateral septum neurotensin neurons to tuberal nucleus controls hedonic feeding. Mol Psychiatry. 2022;27(12):4843–60.

17. Woodworth HL, Beekly BG, Batchelor HM, Bugescu R, Perez-Bonilla P, Schroeder LE, et al. Lateral Hypothalamic Neurotensin Neurons Orchestrate Dual Weight Loss Behaviors via Distinct Mechanisms. Cell Rep. 2017;21(11):3116–28.

18. Kurt G, Kodur N, Quiles CR, Reynolds C, Eagle A, Mayer T, et al. Time to drink: Activating lateral hypothalamic area neurotensin neurons promotes intake of fluid over food in a time-dependent manner. Physiol Behav. 2022;247:113707.

19. Ma C, Zhong P, Liu D, Barger ZK, Zhou L, Chang WC, et al. Sleep Regulation by Neurotensinergic Neurons in a Thalamo-Amygdala Circuit. Neuron. 2019;103(2):323–34 e7.

20. Zhong P, Zhang Z, Barger Z, Ma C, Liu D, Ding X, et al. Control of Non-REM Sleep by Midbrain Neurotensinergic Neurons. Neuron. 2019;104(4):795–809 e6.

21. McHenry JA, Otis JM, Rossi MA, Robinson JE, Kosyk O, Miller NW, et al. Hormonal gain control of a medial preoptic area social reward circuit. Nat Neurosci. 2017;20(3):449–58.

22. Gordon CJ, McMahon B, Richelson E, Padnos B, Katz L. Neurotensin analog NT77 induces regulated hypothermia in the rat. Life Sci. 2003;73(20):2611–23.

23. St-Gelais F, Jomphe C, Trudeau LE. The role of neurotensin in central nervous system pathophysiology: what is the evidence? J Psychiatry Neurosci. 2006;31(4):229–45.

24. Tabarean IV. Neurotensin induces hypothermia by activating both neuronal neurotensin receptor 1 and astrocytic neurotensin receptor 2 in the median preoptic nucleus. Neuropharmacology. 2020;171:108069.

25. Handler CM, Bradley EA, Geller EB, Adler MW. A study of the physiological mechanisms contributing to neurotensin-induced hypothermia. Life Sci. 1994;54(2):95–100.

26. Alexander MJ, Leeman SE. Widespread expression in adult rat forebrain of mRNA encoding high-affinity neurotensin receptor. J Comp Neurol. 1998;402(4):475–500.

27. Nicot A, Rostene W, Berod A. Neurotensin receptor expression in the rat forebrain and midbrain: a combined analysis by in situ hybridization and receptor autoradiography. J Comp Neurol. 1994;341(3):407–19.

28. Hrvatin S, Sun S, Wilcox OF, Yao H, Lavin-Peter AJ, Cicconet M, et al. Neurons that regulate mouse torpor. Nature. 2020;583(7814):115–21.

29. Armstrong WE, Wang L, Li C, Teruyama R. Performance, properties and plasticity of identified oxytocin and vasopressin neurones in vitro. J Neuroendocrinol. 2010;22(5):330–42.

30. Black JA, Vasylyev D, Dib-Hajj SD, Waxman SG. Nav1.9 expression in magnocellular neurosecretory cells of supraoptic nucleus. Exp Neurol. 2014;253:174–9.

31. Frosini M, Valoti M, Sgaragli G. Changes in rectal temperature and ECoG spectral power of sensorimotor cortex elicited in conscious rabbits by i.c.v. injection of GABA, GABA(A) and GABA(B) agonists and antagonists. Br J Pharmacol. 2004;141(1):152–62.

32. Ishiwata T, Saito T, Hasegawa H, Yazawa T, Kotani Y, Otokawa M, et al. Changes of body temperature and thermoregulatory responses of freely moving rats during GABAergic pharmacological stimulation to the preoptic area and anterior hypothalamus in several ambient temperatures. Brain Res. 2005;1048(1-2):32–40.

33. Osaka T. Cold-induced thermogenesis mediated by GABA in the preoptic area of anesthetized rats. Am J Physiol Regul Integr Comp Physiol. 2004;287(2):R306–13.

34. Zhao ZD, Yang WZ, Gao C, Fu X, Zhang W, Zhou Q, et al. A hypothalamic circuit that controls body temperature. Proc Natl Acad Sci U S A. 2017;114(8):2042–7.

35. Osaka T. Prostaglandin E(2) fever mediated by inhibition of the GABAergic transmission in the region immediately adjacent to the organum vasculosum of the lamina terminalis. Pflugers Arch. 2008;456(5):837–46.

36. Tabarean IV, Behrens MM, Bartfai T, Korn H. Prostaglandin E2-increased thermosensitivity of anterior hypothalamic neurons is associated with depressed inhibition. Proc Natl Acad Sci U S A. 2004;101(8):2590–5.

37. Nikolov RP, Yakimova KS. Effects of GABA-transaminase inhibitor Vigabatrin on thermoregulation in rats. Amino Acids. 2011;40(5):1441–5.

38. Meyer-Spasche A, Reed HE, Piggins HD. Neurotensin phase-shifts the firing rate rhythm of neurons in the rat suprachiasmatic nuclei in vitro. Eur J Neurosci. 2002;16(2):339–44.

39. Yamada M, Cho T, Coleman NJ, Yamada M, Richelson E. Regulation of daily rhythm of body temperature by neurotensin receptor in rats. Res Commun Mol Pathol Pharmacol. 1995;87(3):323–32.

40. Yang WZ, Du X, Zhang W, Gao C, Xie H, Xiao Y, et al. Parabrachial neuron types categorically encode thermoregulation variables during heat defense. Sci Adv. 2020;6(36).

41. Tabarean IV. Activation of Preoptic Arginine Vasopressin Neurons Induces Hyperthermia in Male Mice. Endocrinology. 2021;162(2).

42. Paxinos G, Franklin BJ. The mouse brain in stereotaxic coordinates. 2nd ed. San Diego: Academic; 2001.

43. Lundius EG, Sanchez-Alavez M, Ghochani Y, Klaus J, Tabarean IV. Histamine influences body temperature by acting at H1 and H3 receptors on distinct populations of preoptic neurons. J Neurosci. 2010;30(12):4369–81.

